# Temperature scaling of developmental tempo reveals reaction-dominated morphogen patterning

**DOI:** 10.64898/2026.10.06.757014

**Authors:** Daniel Čapek, Theresia Eich, Anna Kögler, Hernán Morales-Navarrete, Oleksandr Riabtsev, Thomas Thumberger, Leonie Floeth, Johanna Bihler, Baubak Bajoghli, Joachim Wittbrodt, Patrick Müller

## Abstract

A central question in developmental biology is how embryonic patterning remains robust when developmental tempo changes. In ectotherms, temperature strongly alters developmental speed and biochemical reaction rates, whereas diffusion is expected to change only weakly. How morphogen gradients required for patterning remain correctly scaled under these conditions is unclear. We addressed this problem in medaka embryos by quantifying developmental speed, morphogen clearance, diffusion, and endogenous Nodal dynamics across a broad temperature range. Developmental tempo, morphogen production, clearance, and tissue movement increased approximately two- to threefold per 10°C, whereas effective morphogen diffusion remained largely unchanged. Despite this mismatch, germ-layer proportions and morphogen patterning remained robust. Our mathematical simulations suggest that this robustness arises because the system operates in a reaction-dominated regime, in which changes in reaction and growth kinetics – rather than diffusion – are sufficient to rescale morphogen gradients. Together, our experiments and simulations provide new insights into how morphogen gradients adjust to maintain accurate tissue patterning across changes in developmental time and tempo.

## Introduction

Embryonic pattern formation relies on gradients of signaling molecules that provide cells with positional information^1^. The shape and amplitude of such morphogen gradients arise from the balance between molecular production, diffusion, and protein degradation^2,3^. While these processes have been extensively characterized under controlled laboratory conditions, natural environments are rarely constant. Most animal species are poikilothermic, and their body temperatures closely follow those of their surroundings, which exposes their embryos to substantial temperature fluctuations across seasons^4^. How such temperature changes affect morphogen transport and the shape and range of signaling gradients remains poorly understood.

According to classical physicochemical theories, temperature should affect the different processes of gradient formation in unequal ways. Biochemical reaction rates typically increase with temperature according to the Arrhenius relationship^5^, which describes an exponential dependence of reaction rate on reciprocal absolute temperature. Over the physiological temperature range, this often corresponds to an approximately two- to threefold change in rate for a 10°C temperature shift, an observation that is known as the Q_10_ rule or van ‘t Hoff relationship^6,7^. Developmental speed in poikilothermic animals – from insects to fish – accordingly scales with temperature in a remarkably consistent manner^8–18^. For example, for the early development of fish embryos we have previously determined Q_10_ values of 2-3 (Supplementary Fig. 1, Supplementary Movie 1, Supplementary Movie 2, Supplementary Movie 3)^11^. By contrast, the Einstein-Stokes relation predicts that diffusion changes only weakly within the physiologically relevant temperature range (*D* ∝ *T*/*η*(*T*))^19^, and a 10°C increase should alter diffusivity by less than ten percent. These differences raise a fundamental question: How can diffusible signaling gradients that provide the spatial information required to pattern the embryo remain appropriately scaled when embryonic time and tempo change?

We addressed this question in the Japanese rice fish medaka (*Oryzias latipes*), an ectothermic vertebrate with exceptional thermal tolerance and experimentally accessible early development. Medaka embryos develop over a wide range of temperatures^20,21^. Their transparency and genetic tractability enable direct visualization of molecular interactions using CRISPR/Cas9-based fluorescent knock-ins^22^. This makes medaka particularly well suited for investigating how morphogen systems respond to changes in developmental temperature. During blastula and gastrula stages, the Nodal ligands Squint and Cyclops, together with their inhibitor Lefty, form a self-regulatory activator–inhibitor reaction-diffusion system that patterns the germ layers^23^. Short-range Nodal signaling promotes mesendodermal fates, whereas the more broadly distributed inhibitor Lefty restricts Nodal signaling and thereby permits ectodermal development^23^. The Nodal/Lefty system therefore provides a tractable model for testing how reaction kinetics, morphogen transport, and tissue dynamics interact to maintain robust patterning when developmental speed changes.

## Results

### Mesendodermal patterning remains robust despite temperature-dependent changes in developmental tempo

We first asked from a theoretical point of view how temperature-dependent reaction and diffusion parameters would be expected to affect a simple morphogen profile. At steady state, a one-dimensional source-degradation gradient can be approximated by

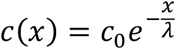

where *c(x)* is the morphogen concentration at distance *x* from the source and *λ* = √(*D*/*k*) is the characteristic length scale defined by the effective diffusion coefficient *D* and the clearance rate constant *k*^3,24^ (Fig. 1a). Because *D* and *k* are expected to scale differently with temperature (Fig. 1b), the model predicts that gradient amplitude, length scale, and threshold positions could become temperature-dependent even if the underlying genetic network is unchanged.

**Figure 1.**
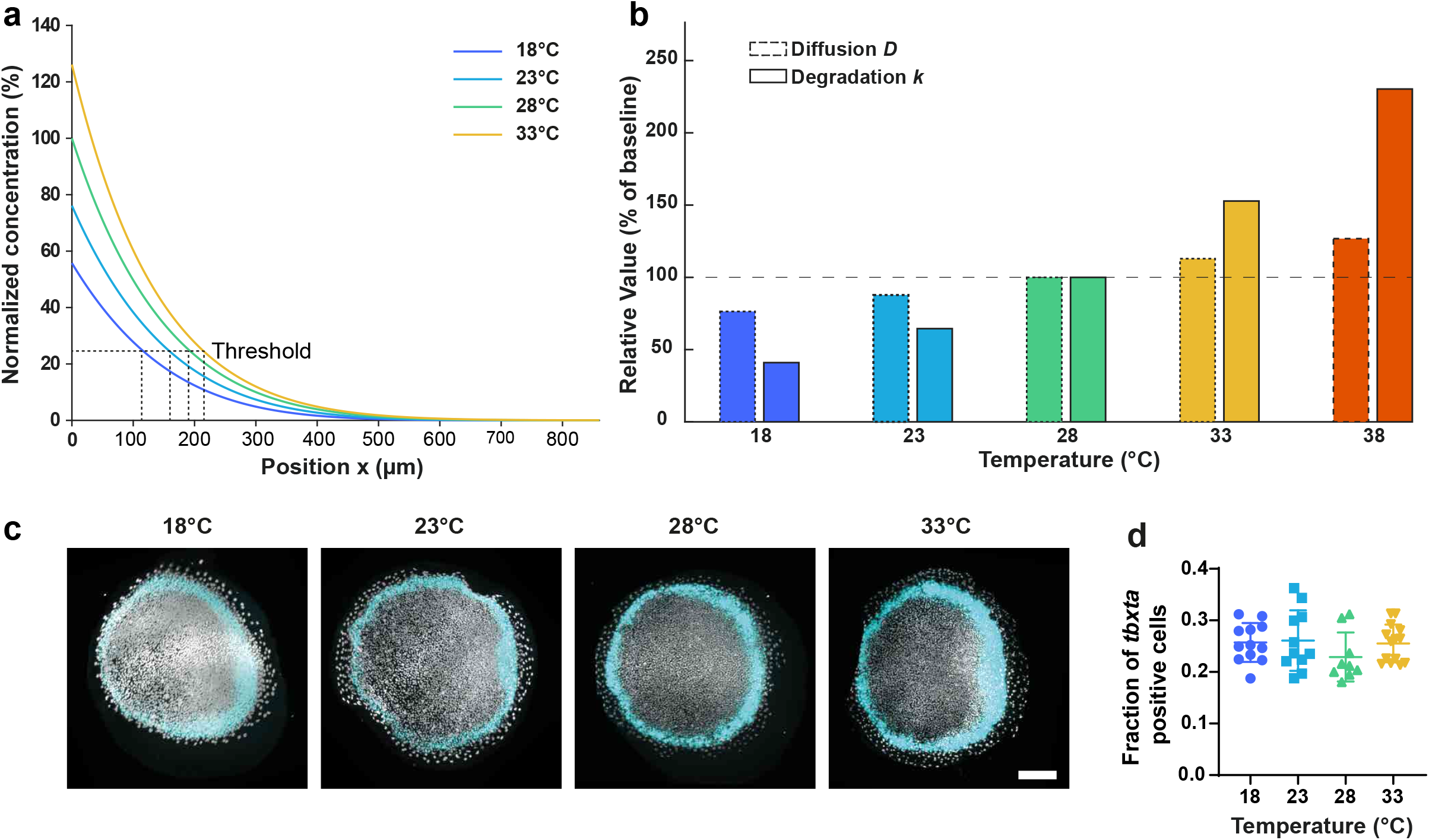
Medaka germ-layer formation is robust across a broad temperature range. **(a)** Hypothetical Nodal morphogen gradients at different temperatures using published zebrafish values for *D* and *k* for 28°C in the classical synthesis-diffusion-degradation model^25^. **(b)** Reaction rates for morphogens such as production and clearance/degradation are expected to scale exponentially with temperature, whereas diffusion should scale linearly according to the Einstein-Stokes equation. **(c)** Fluorescence *in situ* hybridization of medaka embryos raised at different temperature regimes until stage 13. The cyan signal shows cells expressing the mesodermal marker *tbxta*, and the white signal shows nuclei stained with DRAQ7. **(d)** Quantification of (c) showing the fraction of *tbxta*-positive cells. Scale bar: 100 µm.

Using published zebrafish Nodal parameters as a reference^25^ and a flux source with reflecting boundaries, we modeled Q_10_-scaling of reaction rates with weakly temperature-dependent diffusion. However, this simple model did not preserve threshold positions across the full medaka temperature range (Fig. 1a). The simulations therefore predicted that, unless additional compensatory mechanisms exist, germ-layer proportions should be sensitive to developmental temperature.

We tested this prediction by quantifying mesendodermal patterning in medaka embryos raised at 18°C, 23°C, 28°C, or 33°C until the onset of gastrulation (Iwamatsu stage 13^26^). Fluorescent *in situ* hybridization for the Nodal target and mesendodermal marker *tbxta*, combined with DRAQ7 nuclear staining and selective plane illumination microscopy, enabled unbiased quantification of *tbxta*-positive and *tbxta*-negative cells in the animal hemisphere. Despite the large differences in developmental time, the fraction of *tbxta*-positive cells remained stable at approximately 25% across all conditions (24.952% on average; Kruskal-Wallis test, p = 0.3765) (Fig. 1c-d). In contrast, even low doses of chemical Nodal inhibition dramatically reduced *tbxta* expression (Supplementary Movie 4).

To confirm that the temperature treatments altered gastrulation dynamics, we tracked nuclei at the blastoderm margin in embryos expressing H2A-mCherry. Margin cells moved at a velocity of 1.278 µm/min at 18°C and 3.115 µm/min at 28°C, corresponding to an apparent Q_10_ of 2.4 (Supplementary Movie 8). Thus, medaka embryos experience a strong temperature-dependent change in developmental tempo, but still preserve mesendodermal proportions. This robustness suggested that the Nodal/Lefty system compensates for the physical mismatch between reaction-rate and diffusion scaling.

### Morphogen clearance scales with developmental tempo

To investigate how germ-layer proportions remain robust despite changes in developmental tempo, we next quantified the temperature dependence of morphogen clearance. We generated Dendra2-tagged Squint, Cyclops, and Lefty constructs and injected the corresponding mRNAs into one-cell stage medaka embryos. After approximately 7 hours of development, the fusion proteins were pulse-converted, and the converted fluorescence was monitored for 5 h by fluorescence decay after photoconversion (FDAP) assays (Fig. 2a-c, Supplementary Fig. 3, Supplementary Movie 5).

**Figure 2.**
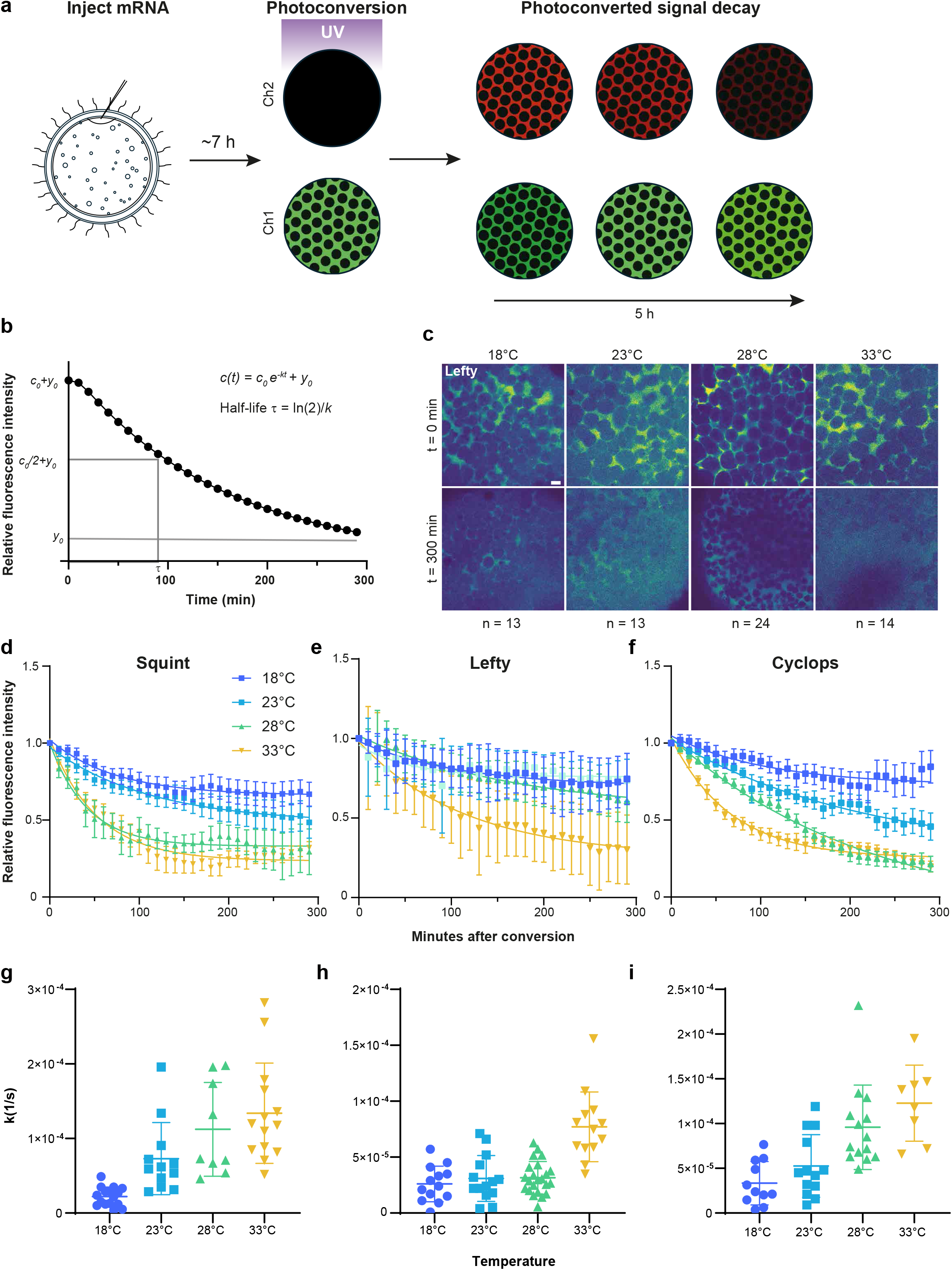
Morphogen stability scales with temperature. **(a,b**) Fluorescence decay after photoconversion (FDAP) principle. mRNA encoding the protein of interest labeled with photoconvertible Dendra2 was injected into one-cell stage medaka embryos, which were then incubated for about 7 h. The green protein (Ch1) was pulse-converted, generating a limited pool of molecules labeled in red (Ch2) and imaged with time-lapse microscopy for 5 h. The half-life and the specific clearance rate of the protein can be determined from the decay of the fluorescence signal (b). **(c)** Representative example fluorescence images of the converted channels of Lefty at different temperatures directly after photoconversion (upper row) and after 5 h (lower row). **(d-f)** Fluorescence decay. Mean normalized fluorescence intensity ± SEM over time for Squint (d), Lefty (e), and Cyclops (f). Solid lines show the exponential decay fit to *I(t*) = *I*_*0*_e^-kt^ + *b*. **(g-i)** Clearance rate constants determined from FDAP experiments. Decay rates (per second) of Squint (g), Lefty (h), and Cyclops (i) at different temperatures. The bars show the mean and the standard deviation, symbols the single values. Scale bar: 20 µm.

Clearance rates increased with temperature for all three proteins (Fig. 2d-i). For Squint, clearance rate constants increased from 0.22 (± 0.13) × 10^−4^/s at 18°C to 0.73 (± 0.49) × 10^−4^/s at 23°C, 1.1 (± 0.62) × 10^−4^/s at 28°C, and 1.3 (± 0.67) × 10^−4^/s at 33°C, corresponding to an average Q_10_ of 2.6. Lefty exhibited overall slower clearance kinetics, with rate constants of 0.26 (± 0.16) × 10^−4^/s at 18°C, 0.31 (± 0.21) × 10^−4^/s at 23°C, 0.32 (± 0.15) × 10^−4^/s at 28°C, and 0.77 (± 0.32) × 10^−4^/s at 33°C, yielding an average apparent Q_10_ of 2.6. For Cyclops, we found clearance rate constants of 0.33 (± 0.19) × 10^−4^/s at 18°C, 0.53 (± 0.35) × 10^−4^/s at 23°C, 0.96 (± 0.47) × 10^−4^/s at 28°C, and 1.23 (± 0.43) × 10^−4^/s at 33°C, resulting in an average apparent Q_10_ value of 2.5. Thus, morphogen clearance scaled with temperature in the same range as whole-embryo developmental tempo^11^ (Supplementary Fig. 1). Fitting the natural logarithm of the clearance rate constants against reciprocal absolute temperature in an Arrhenius analysis yielded an apparent activation energy of 73 kJ/mol, comparable to that reported for early embryonic development^11^.

Interestingly, our measured clearance rates in medaka were close to those previously reported for zebrafish: 1.2 × 10^−4^/s for Cyclops, 1.0 × 10^−4^/s for Squint, and 0.53 and 0.69 × 10^−4^/s for the two Lefty proteins^25^. This similarity is unexpected given the two- to threefold slower development of medaka compared to zebrafish (Supplementary Fig. 1, Supplementary Movie 2, Supplementary Movie 3). Thus, although morphogen clearance scales with temperature within medaka, the comparable clearance rates between medaka and zebrafish argue against protein turnover acting as a general determinant of species-specific developmental tempo in teleost embryos. This finding differs from current models of mammalian development, in which differences in protein stability have been proposed to contribute substantially to species-specific timing in cell culture^27,28^. Together, these observations suggest that the mechanisms controlling developmental tempo may differ between vertebrate lineages.

The strong temperature dependence of morphogen clearance raises the question of how mesendodermal proportions remain stable across temperatures. In a simple diffusion–clearance model, the steady-state gradient length scale is given by *λ* = √(*D*/*k*), where *D* is the diffusion coefficient and *k* is the clearance rate constant^3,24^. The approximately sixfold increase in Nodal clearance between 18°C and 33°C would therefore shorten this length scale by about 2.4-fold if diffusion remained unchanged. Maintaining the same length scale would require a proportional increase in diffusion. We therefore asked whether morphogen diffusion exhibits this compensatory temperature dependence.

### Diffusion is stable across different temperatures and surprisingly slow

To test this possibility, we measured effective diffusion using fluorescence recovery after photobleaching (FRAP) in embryos expressing GFP-tagged Squint or Lefty (Fig. 3a,b, Supplementary Fig. 4, Supplementary Movie 6). Diffusion of the Nodal ligand Squint in medaka embryos was slow across all temperatures, with values of 0.45 (± 0.52) μm^2^/s at 18°C, 0.53 (± 0.33) μm^2^/s at 23°C, 0.34 μm^2^/s (± 0.36) at 28°C and 0.2 μm^2^/s (± 0.1) at 33°C. Lefty diffused much faster than Squint, with values of 11.13 (± 7.9) μm^2^/s at 18°C, 18.41 (± 20.6) μm^2^/s at 23°C, 9.8 μm^2^/s (± 8.4) at 28°C and 9.7 μm^2^/s (± 6.6) at 33°C (Fig. 3c,d).

**Figure 3.**
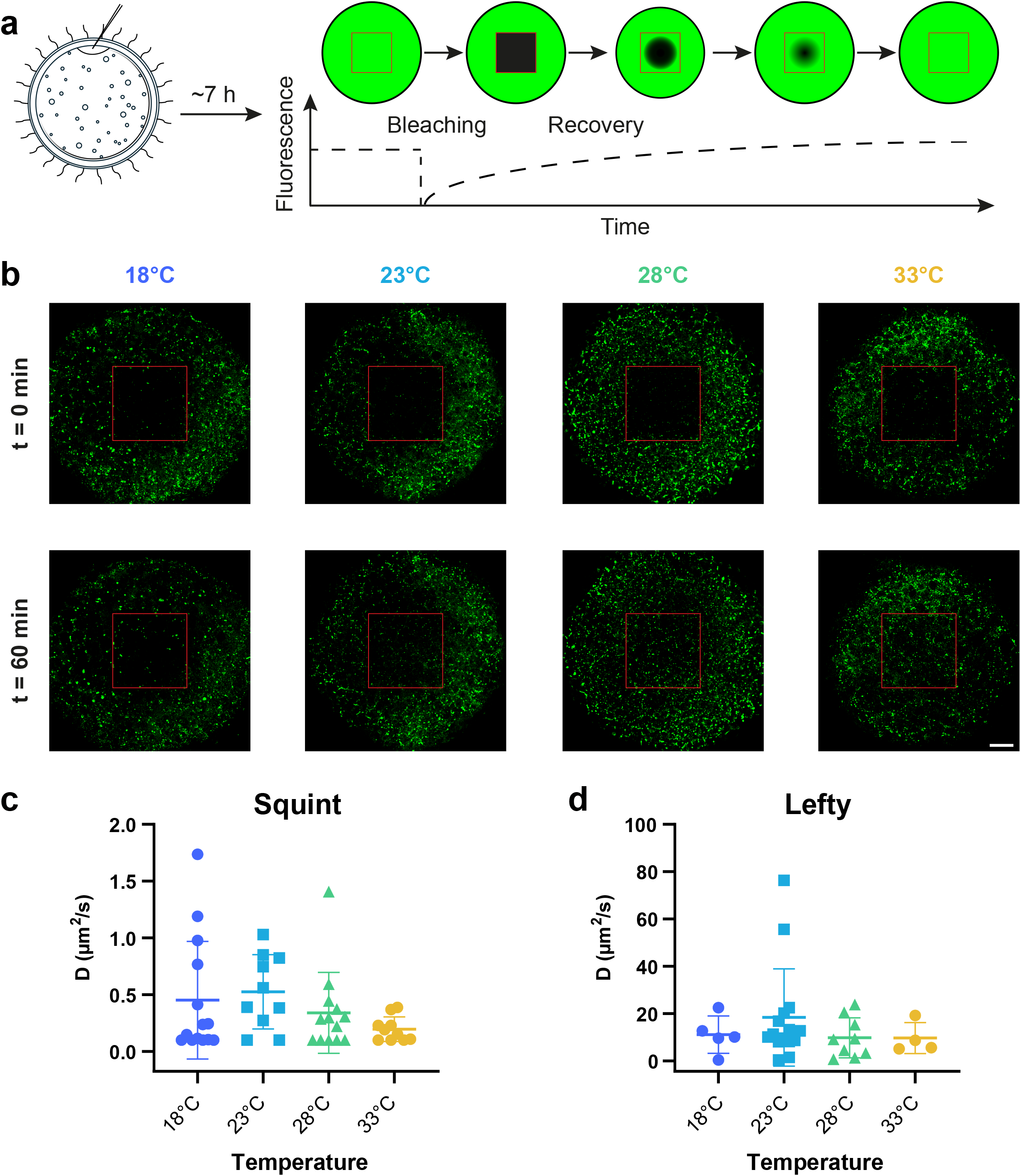
Morphogen diffusion does not scale with temperature. (**a**) Fluorescence recovery after photobleaching (FRAP) principle. Embryos were injected with mRNA encoding GFP-labeled constructs and incubated for about 7 h. One plane of an embryo was imaged with a confocal microscope. A smaller region of interest was bleached by imaging with maximum laser intensity. The recovery of the signal was measured for one hour. **(b)** Representative images of Squint-sfGFP. Examples of Squint recovery at different temperatures directly after bleaching (upper row) and after one hour (lower row). Columns show embryos at 18°C, 23°C, 28°C, and 33°C. Green fluorescence marks Squint-sfGFP. Recovery was recorded at 10 s intervals and fitted to estimate effective tissue-scale diffusion coefficients (representative recovery curves and fits are shown in Supplementary Fig. 4). Scale bar: 50 µm, bleach window (red): 141.7 µm × 141.7 µm. **(c**,**d)** Effective diffusion coefficients. Scatter plots of the diffusion coefficients of Squint (c) and Lefty (d) measured at different temperatures. Each point represents one measurement from a distinct embryo. Bars indicate the mean ± standard deviation.

Thus, the expected differential diffusivity between Nodal and Lefty was preserved^25^, but neither effective Squint nor effective Lefty diffusion increased with developmental temperature. In comparison to zebrafish at 28°C, Lefty showed a similar tissue-scale diffusion coefficient of approximately 10 μm^2^/s, whereas the Nodal ligand Squint was nearly an order of magnitude slower in medaka than zebrafish Nodal (approximately 0.3 μm^2^/s in medaka compared with approximately 3 μm^2^/s in zebrafish^25^). This unusually slow effective Nodal diffusion is consistent with the punctate Squint-GFP distribution in medaka embryonic tissues (Fig. 3b), suggesting that a high fraction of molecules is associated with receptors, cell surfaces, or other tissue structures^29^. These measurements argue against gradient robustness being maintained by coordinated scaling of ligand diffusion and clearance. We therefore considered whether tissue transport and reaction kinetics could reduce the influence of diffusion on patterning.

To compare tissue transport, clearance, and diffusion on the scale of the epiblast, we calculated the Péclet number *Pe* = *vl/D* and the diffusion-based Damköhler number *Da* = *kl*^*2*^*/D*. At 28°C, we used the measured mean margin-cell velocity *v* = 3.115 µm/min (0.0519 µm/s), an initial domain length *l* = 300 µm, the measured Squint diffusivity *D* = 0.34 µm^2^/s, and the measured Squint clearance rate *k* = 1.1 × 10^−4^/s. These values give *Pe* ≈ 46 and *Da* ≈ 29. The corresponding characteristic times are approximately 1.6 h for tissue transport across 300 µm (*l/v*), 2.5 h for clearance (1/*k*), and 74 h for diffusion over the same distance (*l*^*2*^*/D*). Thus, on this spatial scale, both tissue transport and clearance operate much faster than diffusion. For Lefty, whose effective diffusivity at 28°C was much higher (9.8 µm^2^/s), the corresponding Péclet number was only ≈ 1.6.

### Endogenous Squint forms a steep gradient with temperature-dependent dynamics

To relate the exogenous biophysical measurements to endogenous ligand distributions, we generated a homozygous medaka Squint-sfGFP knock-in line by inserting superfolder GFP between the pro-domain and mature domain of the endogenous Squint locus (Supplementary Fig. 2). Light-sheet imaging revealed the onset of detectable Squint-sfGFP expression at stage 11 in the periderm on the dorsal side of the embryo. The signal then spread along the marginal circumference, reaching roughly half of the perimeter by stage 13 (Fig. 4a,b,d). The signal was diffuse at the periderm but strongly enriched in puncta at or near the cell surface in the blastoderm (Supplementary Movie 7). The distribution of Squint-sfGFP largely overlapped with the domain of endogenous *sqt* mRNA expression detected by *in situ* hybridization (Supplementary Fig. 5). At the tissue scale, endogenous Squint-sfGFP formed a steep marginal gradient. Signal intensity peaked close to the margin, extended approximately six cell diameters into the blastoderm, and remained strongly associated with punctate structures at or near cell surfaces (Fig. 4b-d). The peak amplitude increased during early gastrulation, reaching a maximum approximately 3 hours after signal onset at 28°C, and then declined before becoming undetectable by mid-gastrulation (stage 15, Fig. 4d, Supplementary Movie 7). Although low-intensity signal spread further toward the animal pole at later stages, this distal signal was not accompanied by expansion of the *tbxta*-positive mesendodermal domain, consistent with a threshold-based interpretation of Nodal signaling (Fig. 1).

**Figure 4.**
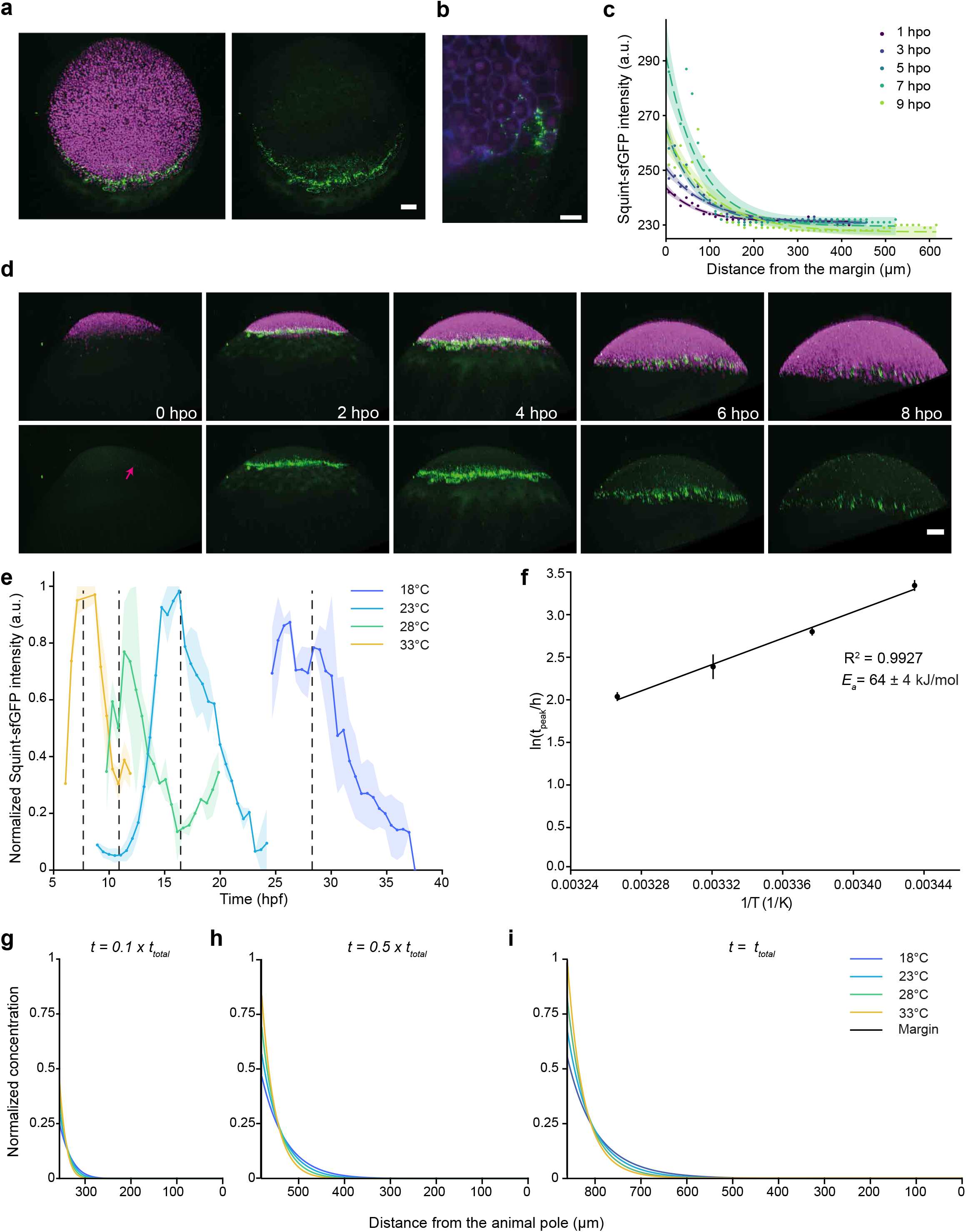
Medaka Squint-sfGFP knock-in and reaction-dominated temperature scaling model of morphogen-mediated patterning. **(a)** Medaka Squint-sfGFP knock-in at stage 13, animal*-*pole view maximum intensity projection with nuclei in magenta and Squint in green. **(b)** Lateral view of Squint (green), nuclei and membranes (magenta) and interstitial fluid labeled (blue), animal pole to the top. **(c)** Squint-sfGFP intensity as a function of distance from the margin in one embryo at 18°C at 1, 3, 5, 7 and 9 hours post-onset (hpo) of the Squint-sfGFP gradient (24.7, 26.8, 28.4, 30.6 and 32.7 hpf). Dots show the median intensity in 13.2 µm bins along the axis from the margin towards the animal pole, dashed lines the fit of *I(x)* = *A*·e^−x/λ^ + *b* and shaded areas the 95% confidence band of the fit. **(d)** Time series at 28°C starting at stage 11 with the onset of Squint signal (magenta arrow). **(e)** Normalized Squint-sfGFP intensity in the gradient region over developmental time at 18°C (blue), 23°C (cyan), 28°C (green), and 33°C (yellow). Lines and dots show the median, shaded areas the median absolute deviation (MAD). Dashed lines, mean peak time at each temperature. n = 3 embryos per time point at 18, 23, and 28°C, and 2–4 at 33°C. **(f)** Arrhenius plot of the Squint-sfGFP peak time (ln(*t*_peak_) versus 1/T). Dots show mean peak time at each temperature. Error bars represent SD on the log scale (n = 2, 3, 3 and 2 embryos at 18, 23, 28 and 33°C). The line indicates the slope of linear regression (R^2^ = 0.993), giving an apparent activation energy of *E*_*a*_ = 64 ± 4 kJ/mol (± SE of the slope). **(g-i)** Moving-boundary model of Nodal transport on a growing epiblast domain, including production at the margin, diffusion, clearance, dilution by tissue growth, and epiboly-associated advection. Reaction-dominated simulations of Nodal gradient formation across 18°C (blue), 23°C (cyan), 28°C (green), and 33°C (yellow) show that gradients remain steep and threshold positions are preserved when production, clearance, and tissue growth scale with developmental tempo while diffusion changes only weakly. See Supplementary Movie 10 for the full dynamic simulation.

To investigate whether Squint-sfGFP punctum velocities vary with temperature, we imaged a small region of the margin at higher magnification, tracked puncta and nuclei, and calculated velocities from frame-to-frame displacements in 3D. We found that puncta move at 0.03 (± 0.0033) μm/s at 23°C and at 0.04 (± 0.0122) μm/s at 33°C on average, while the nuclear velocity was 0.036 (± 0.013) μm/s at 23°C and 0.054 (± 0.0113) μm/s at 33°C (Supplementary Fig. 6a,b). The migration speed measured from nuclear tracking matched our estimated global epiboly rate. Because Squint puncta were closely associated with cells and cell migration was faster at warmer temperatures, we estimated the local nuclear velocity field and subtracted it from puncta trajectories to minimize the influence of advection. This yielded residual puncta velocities of 0.036 (± 0.0038) μm/s at 23°C and 0.044 (± 0.0088) μm/s at 33°C (Supplementary Fig. 6c). Thus, temperature affects the velocity of Squint puncta on a cellular scale, but less strongly than it affects tissue movement.

We next measured the temporal dynamics of endogenous Squint-sfGFP fluorescence across temperatures. At higher temperatures, the signal emerged earlier and persisted for a shorter period (Fig. 4e, Supplementary Movie 9). From these temperature-dependent rates, we determined an apparent activation energy of 64 (± 4) kJ/mol (Fig. 4f), closely matching the values estimated for overall developmental progression and protein clearance.

### Reaction-dominated simulations explain temperature-robust gradient scaling

Together, the slow effective diffusion of Squint and the strong temperature dependence of tissue movement and clearance prompted us to ask whether robust gradient scaling can emerge without diffusion scaling with developmental tempo. To address this question, we developed a minimal one-dimensional model of Nodal transport on a linearly growing domain representing the epiblast during epiboly (Fig. 4g–i, Supplementary Movies 10–14). In the model, ligand is supplied at the moving margin by a flux source, diffuses through the tissue, undergoes first-order clearance, and is diluted as the domain expands uniformly according to *L(t) = L*_*0*_ *+ G*_*t*_. At the reference temperature of 28°C, we used *D* = 0.3 μm^2^/s, *k* = 1.1 × 10^−4^/s, *j* = 1.1 × 10^−4^ mol/(μm^2^·s), and *G* = 3.115 μm/min, consistent with the measured Squint diffusivity, clearance, and epiboly speed.

To simulate development at 18°C, 23°C, 28°C, and 33°C, we scaled production, clearance, and domain growth with the same Arrhenius factor (*E*_*a*_ = 77 kJ/mol), whereas diffusion changed only according to the Einstein-Stokes relation. The total simulation time was scaled inversely with the same Arrhenius factor, allowing comparison at equivalent developmental stages rather than equivalent absolute times. Under these assumptions, the simulated gradients remained steep, and their normalized profiles were largely preserved across temperatures despite only weak changes in *D* (Fig. 4g–i). Thus, robust gradient scaling does not require diffusion to scale with developmental tempo. Instead, when reaction rates and tissue dynamics scale together with developmental speed, morphogen patterning can remain robust in a reaction-dominated regime.

## Discussion

Taken together, our results reveal a striking mismatch in how the processes underlying morphogen patterning respond to temperature. Developmental tempo, tissue movement, and morphogen clearance increase strongly with temperature, whereas morphogen diffusion remains largely unchanged. Despite these differences, germ-layer proportions are remarkably stable across temperatures. This robustness cannot be explained by a simple diffusion-based mechanism in which morphogen range is maintained through coordinated scaling of *D* and *k*. Instead, our results support a reaction-dominated patterning regime in which temperature-sensitive reaction kinetics and tissue dynamics determine the spatial and temporal signaling environment, while diffusion plays a comparatively permissive rather than rate-limiting role. In such a regime, the relevant patterning variable is not the absolute distance that a ligand molecule diffuses per unit clock time, but the balance between ligand production, capture, clearance, signaling duration, and the movement of cells through the signaling field. Because these processes accelerate together with developmental tempo, the signaling history experienced by cells can remain approximately invariant even when absolute time changes. This interpretation is supported by our moving-boundary simulations, in which production, clearance, and tissue expansion scaled with developmental speed while diffusion changed only weakly. This also explains why the endogenous Squint gradient can be steep and membrane-associated without causing temperature-dependent expansion or contraction of the mesendodermal domain.

A recent preprint on medaka somitogenesis similarly showed that robust patterning across temperatures can emerge from the coordinated adjustment of developmental processes with different thermal sensitivities. Despite strongly temperature-dependent oscillation dynamics, segment patterning remained largely temperature-invariant because distinct processes compensated for one another^30^. Together with our findings, these results suggest that temperature robustness does not require all underlying processes to scale identically, but can instead arise from the coordinated interplay of partially compensatory mechanisms that preserve developmental patterning across changing conditions.

The relationship between clearance and developmental tempo also differs within and between species. Despite their markedly different gastrulation tempos, medaka and zebrafish show similar morphogen clearance rates at 28°C. Thus, the clearance rates measured here are unlikely to explain the difference in developmental tempo between these two teleosts, even though their temperature-dependence could contribute to changes in tempo within each species. This contrasts with mouse–human comparisons, in which differences in protein stability are associated with species-specific developmental tempo^27,28^. One possibility is that development in a thermally buffered environment relaxed the constraint to maintain temperature-responsive protein turnover, allowing clearance rates to evolve as species-specific contributors to tempo. Under this hypothesis, the Arrhenius-like temperature dependence observed in fish may reflect an ancestral constraint, whereas its role in setting interspecies differences in mammals emerged later.

Our results provide a physical framework for environmental canalization^16,32,33^ of early vertebrate patterning^31^. Rather than compensating for temperature by tuning diffusion, medaka embryos appear to limit the role of diffusion in setting germ-layer proportions. This strategy differs from expectations derived from reaction-diffusion systems^29,34^ and may represent an adaptation of ectothermic embryogenesis to fluctuating environments. From an evolutionary perspective, this may provide an efficient route to robustness because it does not require extensive rewiring of regulatory networks or developmental programs. Instead, existing molecular networks, tissue dynamics, and physical constraints together foster robustness. More broadly, the findings suggest that robustness of developmental patterning across time and temperature can emerge when reaction networks, tissue dynamics, and morphogen interpretation scale together, while molecular diffusion remains effectively insulated from changes in developmental tempo.

## Acknowledgements

This work was supported by the European Research Council under the European Union’s Horizon 2020 research and innovation programme (grant agreements No. 637840 *QUANTPATTERN*, and No. 863952 *ACE-OF-SPACE*), the Deutsche Forschungsgemeinschaft (DFG, German Research Foundation) under Germany’s Excellence Strategy – EXC 2117 – 422037984, DFG SFB1756 – 550938463, and the Austrian Science Fund (FWF, Erwin Schrödinger Stipendium J 4507-B). We thank the Bioimaging Centre of the University of Konstanz for microscopy support, Manu Čapek, Isabella Kischko and Vilson Karaj for genotyping assistance, and Livia Herse and Rosa Manukjan for assistance with data and file management. We are grateful to NBRP Medaka for providing hatching enzyme.

## Author contributions

Conceptualization, P.M.; Methodology, D.Č., T.E., A.K., H.M.-N., O.R., T.T., J.B., and P.M.;

Investigation, D.Č., T.E., A.K., H.M.-N., O.R., T.T., and J.B.; Formal analysis, D.Č., H.M.-N.,

T.E., O.R., and P.M.; Resources, T.T., B.B., J.W., and P.M.; Writing - original draft, D.Č. and P.M.; Writing - review and editing, all authors; Supervision, P.M.; Funding acquisition, D.Č., P.M.

## Data and code availability

Data generated in this study will be available from https://kondata.uni-konstanz.de after publication. The code for gradient and moving-boundary simulations will be available from https://github.com/mueller-lab after publication.

## Materials and Methods

### Medaka husbandry

Medaka (*Oryzias latipes*) embryos were collected from natural matings and maintained in embryo medium^11,35^ at the indicated temperatures. Embryos were staged according to the classical Iwamatsu staging atlas^26^. Temperature treatments were performed in calibrated incubators or temperature-controlled microscope environments, and experimental temperatures were monitored during incubation and imaging with Sensirion temperature sensors. All animal work was performed in accordance with animal experiment permits 35/185.81-5/EB 03/20 G (Regierungspräsidium Tübingen), 35-9185.81/G-21/096 and 35-9185.81/G-23/092 (Regierungspräsidium Freiburg).

### Generation of fusion constructs

sfGFP-tagged Squint and Cyclops constructs were generated by inserting sfGFP between the pro-domain and mature domain of the corresponding precursor proteins. For Squint, the previously described linker architecture was used^25^. Lefty-sfGFP was generated by adding sfGFP to the C-terminus after a flexible linker^25^ (Supplementary Fig. 2a). Dendra2-tagged constructs for FDAP measurements were generated using the same domain logic, replacing the fluorescent protein with Dendra2. Construct activity was tested by mRNA injection. Nodal constructs were assessed by their ability to broaden expression of the Nodal target gene *goosecoid*^36^, and Lefty constructs by their ability to induce Nodal loss-of-function phenotypes (Supplementary Fig. 2b,c).

### Generation of knock-in lines

Knock-in lines were generated using CRISPR/Cas9-mediated homology-directed repair, following the biotinylated donor strategy described previously^22^. For endogenous Squint-sfGFP, a superfolder GFP tag was inserted between the pro-domain and the mature domain of the *squint* locus (Supplementary Fig. 2d,e). sgRNAs were designed with CCTop as described previously^37^. The genomic *squint* target site was CAAGAGGGAACAACACAGCC (PAM: TGG). The sgRNA spacer sequence was GGAGAGGGAACAACACAGCC, with the first two nucleotides replaced by GG for T7 transcription. Oligonucleotides were ordered from Eurofins, cloned into the DR274 vector, and sgRNAs were transcribed using a MEGAscript T7 Transcription Kit (Invitrogen AM1334). The donor construct contained homology flanks and silently mutated target sites to prevent re-cutting after integration. The donor sequence was amplified from a pCS2 vector using the biotinylated primers G*C*T*G*C*CAGCAGGTTGGTAA and C*A*T*G*A*A*TGCATGGTTAGTTGGTCT.

Embryos were injected at the one-cell stage with heiCas9-encoding mRNA^38^ (150 ng/μl), locus-specific sgRNAs (5 ng/μl), and the biotinylated donor (15 ng/μl). Candidate founders were screened for fluorescence and by genotyping, and precise integration of the *squint-sfGFP* locus was confirmed by PCR and Sanger sequencing in F2 fish (Supplementary Fig. 2f). Homozygous embryos were identified by genotyping and imaged by light-sheet microscopy from stage 11 through gastrulation.

### Fluorescent in situ hybridization and cell quantification

Antisense DIG-labeled RNA probes against *tbxta* and *squint* were generated from template DNA^39^ by *in vitro* transcription. Embryos were fixed at Iwamatsu stage 11 or 13 in 4% ice-cold formaldehyde overnight at 4°C with mild rocking, washed in PBS and dechorionated manually, washed in 100% methanol, and then stored in methanol at −20°C. Fluorescent *in situ* hybridization was performed as described previously^40^, using fluorescein for signal detection. Briefly, rehydrated embryos were hybridized at 65°C overnight with the respective probes and after stringency washes with decreasing formamide and salt levels were blocked for at least 2 h at room temperature in 2% blocking reagent (Roche) in maleic acid buffer containing 0.1% Tween-20. Hybridized probes were detected using HRP-conjugated anti-DIG Fab fragments (Roche; 1:1000) overnight at 4°C. Following washes in PBS containing Tween-20, signals were developed using the TSA Plus Fluorescein kit (PerkinElmer; 1:75 in amplification buffer) for 90 minutes at room temperature in the dark. Embryos were washed, counterstained with DRAQ7 (1:1000 Invitrogen D15106), and imaged on a Lightsheet Z.1 microscope (Zeiss, ZEN black). Embryos that became distorted during fixation were excluded before analysis. Cell numbers in the animal hemisphere were quantified in three dimensions using Imaris 7 (Bitplane). A volumetric mask was generated from the *tbxta* fluorescence signal to delineate the expression domain. The DRAQ7 channel was duplicated, and voxel intensities outside this mask were set to zero in the duplicate channel. Nuclei were then segmented and counted separately in the original and masked DRAQ7 channels using the built-in Spot tool, yielding the total number of nuclei and the number within the *tbxta*-positive domain, respectively. The fraction of *tbxta*-positive cells was calculated as the ratio of these counts for each embryo and compared across temperatures.

### Chorion removal

The choria of living embryos at the four-cell stage onward were mechanically abraded by rolling the eggs over sandpaper and washing 3 times for 5 min each in Milli-Q H_2_O in a 48-well plate. Eggs were then incubated in hatching enzyme for 20 min to 1.5 h. Remaining chorion material was removed with forceps, and dechorionated embryos were transferred into 1x balanced salt solution (BSS: 111 mM NaCl, 5.4 mM KCl, 0.81 mM MgSO_4_, 1.36 mM CaCl_2_, 15 mM HEPES, pH 7.0^41^).

### FDAP

Fluorescence decay after photoconversion (FDAP) was performed on a Zeiss LSM 700 confocal microscope equipped with a 25× objective (LD LCI-PlanApochromat Multi immersion 0.8 NA). mRNAs encoding Dendra2-tagged Squint, Cyclops, or Lefty were injected into one-cell stage embryos. After approximately 7 h of development, embryos were mounted in 1.5% low-melting point agarose on 35 mm µ-Dishes (Ibidi 81158) and covered with embryo rearing medium^42^ (ERM: 17 mM NaCl, 0.4 mM KCl, 0.27 mM CaCl_2_, 0.65 mM MgSO_4_). Multiple embryos were first recorded in both the green and the red channel using a 488 nm and 555 nm diode laser to acquire a pre-conversion image. Green emission was detected through an SP 555 filter (up to 550 nm), whereas red emission was separated by the variable secondary dichroic beamsplitter and detected up to 620 nm. Dendra2 was photoconverted with ultraviolet illumination by a 405 nm diode laser (3% laser power for 1 minute), and converted fluorescence was measured every 10 min for 5 h. Background-corrected intensity traces were normalized to the first post-conversion time point and fitted with the single-exponential decay model *I(t*) = *I*_*0*_e^-kt^ + b, to estimate half-lives and clearance rate constants^43,44^. Temperature was controlled by a stage-top incubation chamber, and for 18°C and 23°C the room temperature was cooled to 15°C first.

To control for imaging-induced bleaching, Squint-Dendra2 embryos were imaged directly after photoconversion for 150 frames at 10 s intervals. The signal remained stable except for small fluctuations due to cell movement (Supplementary Fig. 3).

### FRAP

FRAP measurements were performed on a Zeiss LSM880 system equipped with a 25× (LD LCI-PlanApochromat Multi immersion 0.8 NA) objective. Embryos expressing sfGFP-tagged Squint or Lefty were mounted in 1.5% agarose on 35 mm µ-Dishes (Ibidi 81158), covered with ERM, and imaged with a 488 nm argon laser at 5% intensity to acquire a pre-bleaching image (nominal emission range 483-557 nm). A defined region of interest was photobleached with maximum laser intensity, and fluorescence recovery was monitored for 60 min with 10 s intervals under the same conditions as the pre-bleaching image. Recovery curves were fitted to determine tissue-scale diffusion coefficients^45^. Temperature was controlled by a stage-top incubation chamber, and for 18°C and 23°C the room temperature was cooled to 15°C first.

### Chemical inhibition of Nodal signaling

To inhibit Nodal signaling in the *tbxta:Venus* line^46^, embryos were incubated in SB-505124 hydrochloride hydrate (Merck S4696) at either 2 or 3 µM in medaka ERM starting at stage 8. The drug was left in the medium for the whole time of the experiment.

### Imaging of knock-in lines

Embryos were mounted in 1.5% low-melting-point agarose in 1× BSS in glass capillaries and imaged on a Lightsheet Z.1 microscope (Zeiss, ZEN black) with a 10×/0.5 water-immersion detection objective (zoom 0.7). The chamber was filled with 1× BSS and kept at the target temperature by the microscope’s temperature control. Nuclei were labeled by injection of H2A-mCherry^47^ mRNA (50 ng/µl), membranes with membrane-RFP-encoding mRNA^25^ (100 ng/µl) at the 1-cell stage. Interstitial fluid was labeled by injection with 0.5 ng of Dextran-Alexa Fluor647 (Thermo D22914) into the extracellular space after dechorionation. Squint-sfGFP was excited at 488 nm and detected through a 505–545-nm band-pass filter (exposure 110 ms). z-stacks (pixel size 0.662 µm, z-step 1.87 µm) were acquired every 31.6–32.2 min, starting at 6– 10 hpf (23, 28 and 33°C) or at 24.7 hpf (18°C). Developmental time was calculated from the acquisition time stamps of the CZI files, with fertilization set to the time of egg collection (08:30).

Maximum-intensity projections (MIPs) of the 488-nm channel were calculated from the CZI files for every time point. In Fiji^48^, a polygonal region of interest (ROI) around the Squint-sfGFP signal at the margin was drawn by hand for each time point, and one ROI in the yolk was drawn per embryo. For each time point, the median intensity of the yolk ROI (background) was subtracted from the mean intensity of the Squint-sfGFP ROI. At 18 and 23°C, time points without a localized Squint-sfGFP signal (no ROI) were set to zero, and the 18°C curve was truncated at 38 hpf. At 28 and 33°C, the curves were truncated at 20 hpf and 12 hpf, respectively. Embryos from different recordings were matched to the nearest time point (tolerance 0.3 h), and the median and MAD across embryos were calculated for each time point.

The peak time (t_peak_) of each embryo was obtained by fitting a Gaussian with offset, *y(t)* = *y*_*0*_ + *a*·e^(−(*t* − *tpea*k)2/(2*Σ*2))^, to its normalized curve. Embryos whose maximum was the first recorded time point were excluded because the peak had occurred before the recording started (one embryo at 18°C and two at 33°C). The temperature dependence was described by the Arrhenius equation, *k* = *A*·e^(−Ea/RT)^, with *k* ∝ 1/*t*_*peak*_, so that ln(*t*_*peak*_) = *E*_*a*_*/RT* + *c*. The logarithm of the mean *t*_*peak*_ at each temperature was plotted against 1/*T* (*T* in K), and *E*_*a*_ was calculated from the slope of a linear least-squares fit (*E*_*a*_ = slope × *R, R* = 8.314 J/(mol·K)). Error bars show the SD of *t*_*peak*_ on the log scale.

For gradient analysis, the z-stack of each hourly time point was binned 2 × 2 in xy, resampled in z to isotropic 1.32-µm voxels, rotated by 90° in the image plane and by 45° about the vertical image axis, and maximum-projected. In Fiji^48^, a quadrilateral ROI reaching from the margin to the animal pole was drawn for each time point. The two corners at the animal pole were kept fixed, and the margin-side edge was set to a constant width of 116 µm. Pixels inside the ROI were projected onto the axis from the midpoint of the margin-side edge (0 µm) to the midpoint of the animal-pole edge, and the median of the raw pixel values was calculated in 10-pixel (13.2-µm) bins across the full ROI width. *I(x)* = *b* + *A*·e^(−x/λ)^ was fitted to the bin medians by non-linear least squares, and the 95% confidence band was calculated from the parameter covariance (delta method). The decay lengths were λ = 72 ± 10, 94 ± 11, 67 ± 8, 75 ± 13, 102 ± 18 µm (fit ± SE) at 1, 3, 5, 7 and 9 hpo. ROIs were drawn in Fiji (ImageJ 2.18.0/1.54p). Image analysis, curve fitting and statistics were performed in Python (v3.10) with NumPy (v1.26), SciPy (v1.14), pandas, matplotlib (v3.10), tifffile, roifile and aicspylibczi.

### Epiboly measurements, puncta and tissue-motion analysis

Medaka embryos were imaged every 2 minutes using a 10× objective (W Plan-Apochromat, 0.5 NA, M27 75mm) for whole embryo imaging or every 30 seconds using a 20× objective (W Plan-Apochromat 1.0 NA DIC M27 70mm) for Squint puncta tracking. For nuclear tracking, embryos were injected at the one-cell stage with mRNA encoding H2A-mCherry^47^ and imaged during epiboly.

For image processing and nuclear segmentation, three-dimensional image stacks were first rescaled by a factor of 0.5 in the xy plane and interpolated to isotropic voxel dimensions. Images were intensity-normalized, and the nuclear channel was segmented using Cellpose in 3D^49^. Before segmentation, volumes were downsampled fourfold in each spatial dimension and normalized by clipping intensities to the 10^th^ and 99.9^th^ percentiles. Cellpose was run in 3D with a flow threshold of 0.8 and a cell-probability threshold of 0. The resulting instance-label volumes were resized to the dimensions of the processed image stacks using nearest-neighbor interpolation to preserve discrete labels.

Nuclei and Squint-sfGFP puncta were subsequently tracked in three dimensions using TrackMate v8.1.6 in Fiji^48^. Track coordinates were exported in physical units (µm) together with their corresponding time points. The time interval between consecutive image frames was 30 s. For each trajectory, positions were ordered by time and three-dimensional displacement vectors were calculated between consecutive observations as Δ**r**_i_ = **r**_i_ − **r**_i−1_. Instantaneous velocity vectors were calculated by backward finite differences, **v**_i_ = Δ**r**_i_/Δt_i_, where Δt_i_ = t_i_ − t_i−1_. Instantaneous speed was defined as the Euclidean magnitude of the velocity vector, |**v**_i_| = √(v_x_^2^ + v_γ_^2^ + v_z^2^). Thus, when a trajectory contained a gap of more than one imaging interval, the displacement was divided by the actual elapsed time rather than assuming a single-frame interval.

High-magnification light-sheet movies of Squint-sfGFP embryos were used to track fluorescent puncta and nuclei. To distinguish punctum movement relative to the surrounding tissue from displacement caused by bulk tissue motion, a local three-dimensional nuclear velocity field was estimated independently for each embryo. For every punctum position and time point, the local tissue velocity was calculated from the six spatially nearest tracked nuclei by inverse-distance weighting. For a punctum at position **x**, the nuclear velocity field was estimated as

v_tissue(x) = Σ_j_ w_j_v_j_ / Σ_j_ w_j_, with weights 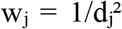, where d_j_ is the three-dimensional Euclidean distance between the punctum and nucleus j. If nuclear velocities were available at the exact punctum time point, spatial interpolation was performed directly at that time. When the punctum time fell between two available nuclear time points, the velocity field was first spatially interpolated independently at the preceding and following time points and then linearly interpolated in time. If data were available on only one side of the queried time point, the closest available nuclear velocity field was used.

Punctum motion relative to the tissue was then calculated by subtracting the interpolated local nuclear velocity vector from the measured punctum velocity vector,

v_rel = v_punctum − v_tissue,

and relative punctum speed was defined as |v_rel| = √(v_rel,x^2^ + v_rel,y^2^ + v_rel,z^2^). Tissue-motion-corrected punctum trajectories were reconstructed by numerical integration of these relative velocity vectors, using the original position of the first tracked punctum as the trajectory origin and updating subsequent positions as r_i_,corr = r_i−1_,corr + v_i_,rel Δt_i_.

For quantitative comparisons, instantaneous speeds were averaged within each trajectory to obtain a mean speed per track. Track-level means were subsequently averaged within each embryo to obtain an embryo-level mean, thereby preventing embryos containing larger numbers of tracked objects from contributing disproportionately to group-level estimates. Temperature-dependent summaries were calculated from these embryo-level values.

### Mathematical modeling of morphogen gradient formation

One-dimensional gradients were simulated using a synthesis-diffusion-clearance model with reflecting boundaries. Baseline diffusion and clearance parameters were based on published zebrafish Nodal measurements^25^. Temperature-dependent reaction rates were scaled using the Arrhenius relation^5^, and diffusion was scaled according to the Einstein-Stokes relation with viscosity correction^19,25,29^.

For the moving-boundary simulations, we modeled Nodal transport and clearance on a linearly growing one-dimensional domain representing the epiblast during epiboly. The domain length was *L(t)* = *L*_*0*_ *+ G*_*t*_ with L_0_ = 300 μm, and ligand concentration *c(x,t)* obeyed ∂*c/∂t + ∂(vc)/∂x = D∂*^*2*^*c/∂x*^*2*^ *-kc*, with *v(x,t) = Gx/L(t)*. The left boundary was sealed, and ligand entered through a flux source *j* at the moving margin. Transforming to the fixed coordinate *ξ = x/L(t)* yielded *∂c/∂t = D/L(t)*^*2*^ *∂*^*2*^*c/∂ξ*^*2*^ *-(k + G/L(t))c*, with *∂c/∂ξ* = 0 at ξ = 0 and *∂c/∂ξ = jL(t)/D* at *ξ* = 1. At the reference temperature of 28°C, *D*_*ref*_ = 0.3 μm^2^/s, *k*_*ref*_ = 1.1 × 10^−4^/s, *j*_*ref*_ = 1.1 × 10^−4^ mol/(μm^2^· s), and *G*_*ref*_ = 3.115 μm/min. *D(T)* was calculated from the Einstein-Stokes relation using water viscosity, while *k(T), j(T)*, and *G(T)* were scaled with a common Arrhenius factor using *E*_*a*_ = 77 kJ/mol. The total integration time was scaled inversely to compare equivalent biological intervals, with *t*_*ref*_ = 10,800 s at 28°C. The transformed PDE was solved in MATLAB using pdepe on *ξ* ∈ [0,1] with a uniform spatial grid and temperature-specific integration times (relative and absolute error tolerances of 10^-3^ and 10^-6^, respectively). For comparative scaling, half-lives were calculated as ln(2)/*k* and static diffusion–clearance lengths as √(*D*/*k*). Péclet and diffusion-based Damköhler numbers were calculated as Pe = v*l*/D and Da = k*l*^2^/D, respectively, with *l* = 300 μm and v converted to μm/s. Experimental mean diffusivities were used for these comparisons. The temperature scaling, coordinate transformation, and numerical implementation are described in detail below.

Diffusivity was assumed to follow the Einstein–Stokes relation

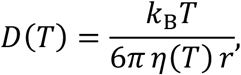

where the dynamic viscosity of the medium *η*(*T*) is assumed to be similar to water, and the particle radius *r* is extracted from the reference value *D*_ref_ via

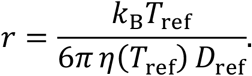

The remaining three rates are scaled with temperature using the Arrhenius law, assuming a common activation energy *E*_a_ = 77 kJ/mol determined for medaka development^11^:

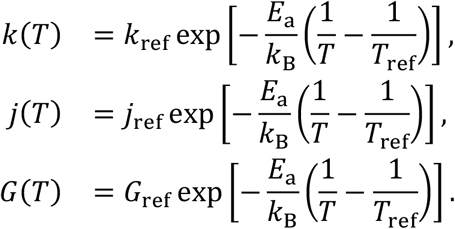

To compare the same biological interval across temperatures, the total integration time is scaled inversely:

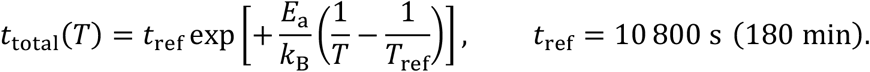

Here, k_B_ is the Boltzmann constant, and temperatures are expressed in Kelvin. We make the following modeling assumptions:

The tissue domain grows linearly in time from its initial length *L*_0_ = 300 *µ*m:

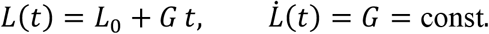

Morphogen is secreted with constant inward flux *j* at the moving right boundary *x* = *L*(*t*) (the advancing epiboly margin); the left boundary *x* = 0 is sealed (zero flux).

The morphogen concentration *c*(*x, t*) evolves according to the parabolic equation

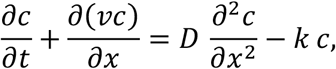

where 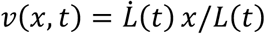 is the tissue velocity field due to uniform stretching. The initial condition is *c*(*x*, 0) = 0 throughout the domain.

The moving boundary makes direct numerical solution inconvenient. We therefore introduce the normalized coordinate

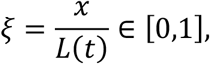

which maps the growing domain onto the fixed interval [0,1] at all times. The domain length grows linearly from its initial value: *L*(*t*) = *L*_0_ + *G t* with constant speed 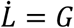.

To express the PDE in terms of ξ we need the transformation rules for the spatial and temporal derivatives. From the definition of ξ we have *x* = ξ*L*(*t*); holding *t* fixed gives

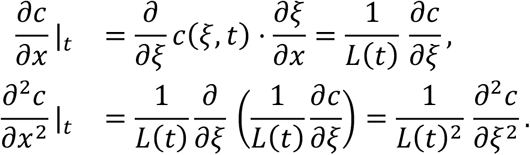

The time derivative at a fixed physical position *x* is more involved because ξ changes in time. Applying the chain rule,

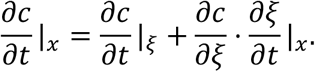

The partial derivative of ξ with respect to *t* at constant *x* is

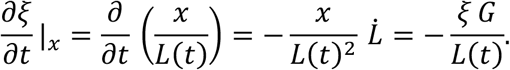

Substituting the expression for the coordinate derivative into the temporal chain rule yields

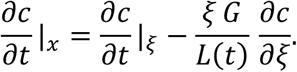

The governing PDE in physical coordinates, after expanding the advection term ∂(*vc*)/ ∂*x* and using *v* = *Gx*/*L*(*t*), reads

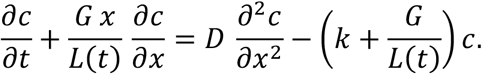

Now substitute the transformed derivatives derived above into the expanded governing PDE. The time derivative becomes ∂*c*/ ∂*t*|_ξ_ − (ξ*G*/*L*) ∂*c*/ ∂ξ. The advection term on the left is

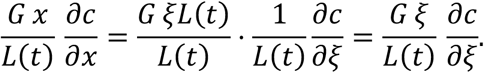

Adding the transformed time derivative and the advection term gives

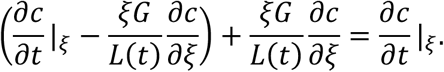

The two ξ*G*/*L* terms cancel exactly, removing the explicit advection. The right-hand side of the expanded governing PDE becomes

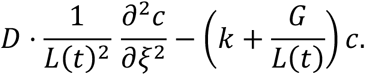

Thus, the transformed PDE on the fixed domain ξ ∈ [0,1] is

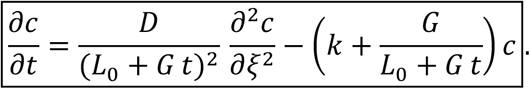

Using ∂*c*/ ∂*x* = (1/*L*) ∂*c*/ ∂ξ, the physical conditions become:

*Left boundary (*ξ = 0)

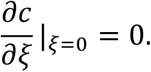

*Right boundary (*ξ = 1)

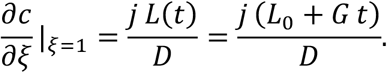

The simulations were implemented in MATLAB and are organized into four components. The core directory contains six files implementing the physics: runSimulation.m is the solver kernel, pdefunGrow.m and bcfunGrow.m supply the PDE coefficients and boundary conditions, buildArrheniusParams.m assembles the temperature-dependent parameter struct, and domainLength.m and waterViscosity.m are utilities for the domain geometry and fluid viscosity respectively. Validation scripts are collected in tests/, figures are produced by main.m, and animations by animate.m.

The transformed PDE above was solved on the fixed interval ξ∈[0,1] using MATLAB’s built-in pdepe solver, which targets parabolic–elliptic equations. The solver accepts six required inputs: a symmetry flag *m*, a PDE coefficient function, an initial-condition function, a boundary-condition function, and the spatial and temporal grids. The flag *m* selects the coordinate geometry: *m* = 0 for Cartesian. The core call inside wrapper runSimulation.m is:

~~~
sol = pdepe(0, …
@(x,t,u,dudx) pdefunGrow(x, t, u, dudx, …
params.D, params.k, params.G, … params.L0), …
@(~) ic_value, …
@(xl,ul,xr,ur,t) bcfunGrow(xl, ul, xr, ur, t, …
params.j, params.G, …
params.L0), …
x_grid, t_grid);
~~~

The spatial and temporal grids are both uniform, constructed as:

~~~
x_grid = linspace(0, 1, N_x);
t_grid = linspace(0, params.t_total, N_t);
~~~

where *N*_*x*_ = 1000 and *N*_*t*_ = *t*_total_*10, giving an output interval of approximately 0.1 s. Because pdepe is required to return a solution at every point in t_grid, this also serves as an upper bound on the internal step size of the ode15s integrator.

The solver accepts equations that can be represented in a form

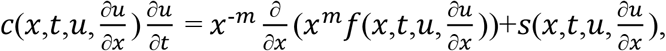

where *m*∈{0,1,2} selects slab, cylindrical, or spherical symmetry respectively. Here *x* is the spatial coordinate, *t* is time, *u*(*x,t*) is the unknown solution field, and ∂*u*/∂*x* is its spatial derivative. The three coefficient functions are supplied by the user as a single function handle and have the following general roles. *c*(*x,t,u*,∂*u*/∂*x*) is the capacity matrix, and its elements couple the partial derivatives with respect to time. *f*(*x,t,u*,∂*u*/∂*x*) is the flux; pdepe places it inside the divergence operator, so every term involving a spatial derivative of *u* is encoded here. *s*(*x,t,u*,∂*u*/∂*x*) is the source, which collects all remaining terms that do not appear inside a spatial derivative, i.e. reaction, production, and clearance terms.

To map the transformed PDE above onto this form, we identify *x* ≡ ξ, *u* ≡ *c*, and *m* = 0, and read off the three coefficients by inspection:

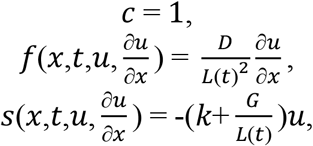

The initial condition is supplied as a scalar ic_value = 0, meaning *u*(*x*,0) = 0 throughout the domain, reflecting the assumption that no morphogen is present at the onset of epiboly.

The boundary conditions are supplied to pdepe via a function handle with signature

[pl, ql, pr, qr] = bcfun(xl, ul, xr, ur, t)

where xl, ul and xr, ur are the spatial coordinate and solution value at the left and right boundary respectively, and *t* is the current time. The boundary points are always the first and last points of x_grid, so xl=0 and xr=1. At these points pdepe does not apply the PDE; instead it enforces the constraint

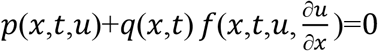

by finding the value of *u* at each boundary that satisfies it, where *f* is the same flux returned by pdefunGrow. The role of *p* and *q* is to encode the physical boundary condition in this form: one starts from the required condition on *f* and rearranges it into *p*+*q*⋅*f*=0 to read off *p* and *q*.

At the left boundary the physical condition is zero flux, *f*|_*x*=0_ = 0. This is already in the form of the equation above with *p*_1_ =0 and *q*_1_ =1. Substituting 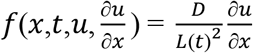 makes it explicit:

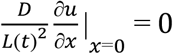

At the right boundary the physical condition is *f*|_*x*=1_=*j*/*L*(*t*), which rearranges to −*j*/*L*(*t*)+*f*=0, giving *p*_*r*_=-*j*/*L*(*t*) and *q*_*r*_=1. Substituting the flux coefficient yields

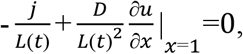

which rearranges to

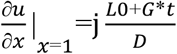

The implementation in bcfunGrow follows directly: the spatial and solution arguments are discarded since neither *p* nor *q* depend on *x* or *u* in this model, and *L*(*t*) is recomputed from *G* and *L*_0_ at the current time *t*.

~~~
function [pl, ql, pr, qr] = bcfunGrow(~, ~, ~, ~, t, j, G, L0) Lt = L0 + G * t;
pl = 0; ql = 1;
pr = −j / Lt;
qr = 1;
end
~~~

### Statistics

Outlier removal for FDAP used the ROUT test with Q = 0.5%. Unless stated otherwise, data are shown as mean ± standard deviation. Kruskal–Wallis tests compared embryo-level *tbxta*-positive cell fractions across temperatures. The epiboly comparison comprised one embryo at 18°C and one at 28°C. Half-lives and characteristic length scales reported in the interspecies comparison were calculated from mean rate constants and diffusivities.

## Supplementary Figure Legends

**Supplementary Figure 1.**
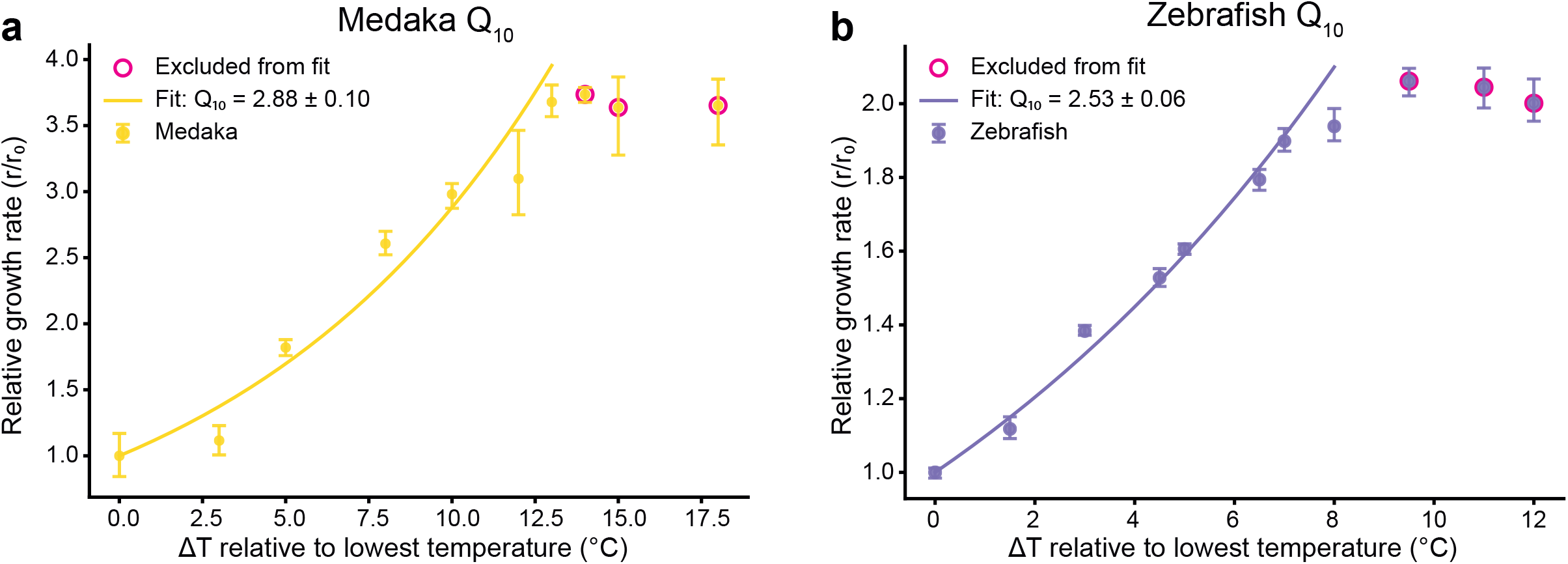
Temperature scaling and Q_10_ values. **(a)** Developmental speed in medaka embryos was quantified across temperatures using high-throughput morphometric staging data^11^, yielding a Q_10_ value of 2.88. Fit and estimated Q_10_ factor for medaka development starting at 18°C. n(18°C) = 65, n(21°C) = 32, n(23°C) = 92, n(26°C) = 47, n(28°C) = 46, n(30°C) = 41, n(31°C) = 21, n(32°C) = 40, n(33°C) = 42, n(36°C) = 35. **(b)** Fit and estimated Q_10_ factor for zebrafish development starting at 23.5°C. n(23.5°C) = 211, n(25°C) = 198, n(26.5°C) = 209, n(28°C) = 168, n(28.5°C) = 126, n(30°C) = 187, n(30.5°C) = 102, n(31.5°C) = 130, n(33°C) = 98, n(34.5°C) = 70, n(35.5°C) = 119. Developmental rates in the plateauing temperature regime were excluded from the fit (magenta circles). Error bars represent 99.99% confidence intervals from bootstrapping with 100 repetitions around the estimated slope of the linear fit to the data. Q_10_ values were estimated from our previously published developmental rates at different temperatures^11^. For comparison, epiboly speed changed from 1.278 μm/min at 18°C to 3.115 μm/min at 28°C, corresponding to an apparent Q_10_ of 2.43. FDAP-derived clearance rates for Squint, Lefty, and Cyclops yielded average apparent Q_10_ values of 2.63, 2.61, and 2.47, respectively, whereas diffusion showed only weak temperature dependence.

**Supplementary Figure 2.**
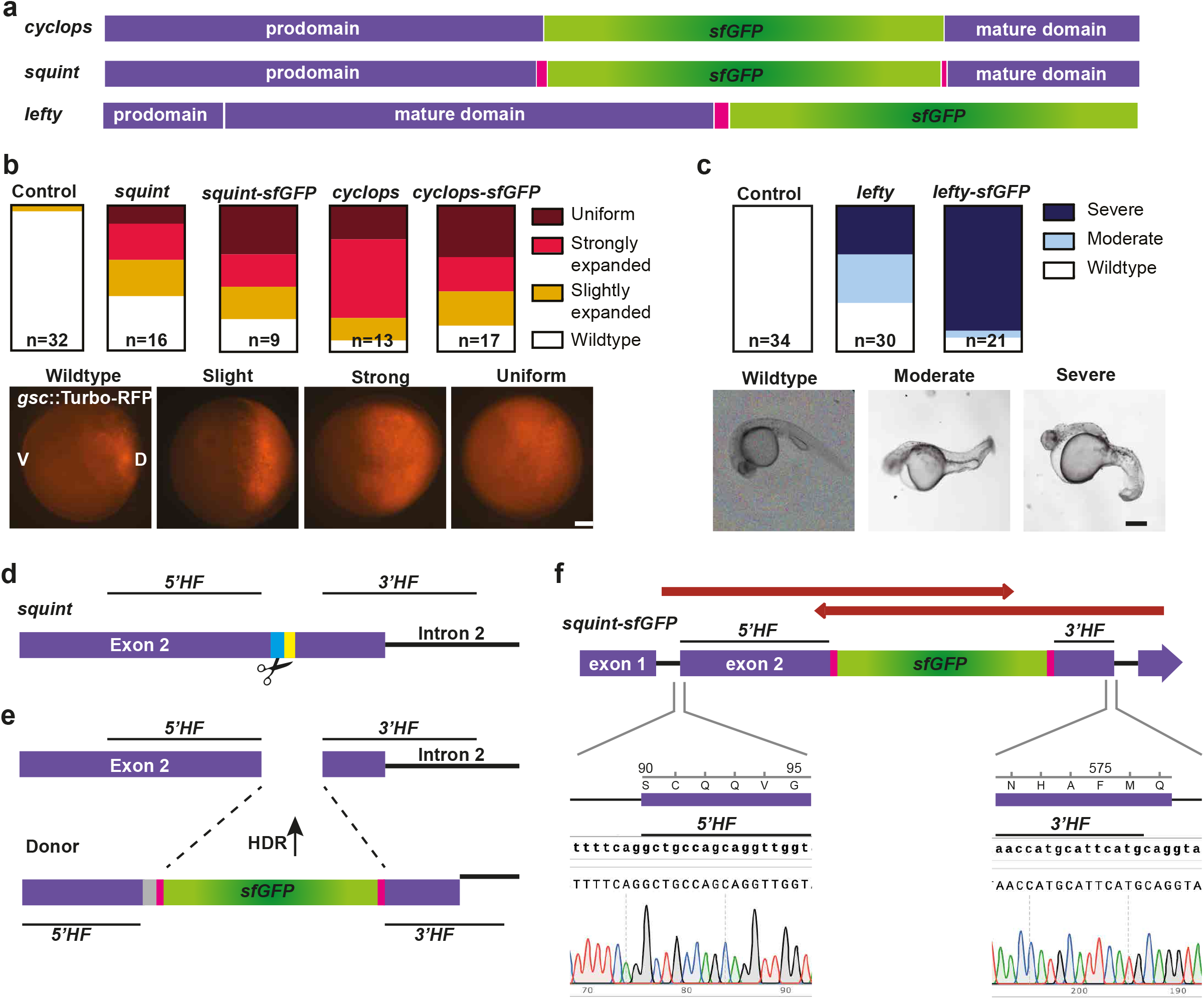
Construct activity and CRISPR/Cas9 knock-in. **(a)** Schematic of the fusion constructs. The sfGFP-encoding sequence was inserted between the pro- and mature domains of Cyclops and Squint, respectively. In the case of Squint, linker sequences (magenta) were used as shown before^25^. For Lefty, sfGFP was added at the C-terminus after a linker sequence^25^. **(b)** The functionality of the Cyclops and Squint constructs was determined by their ability to broaden the expression of the Nodal target gene *goosecoid* (*gsc*^36^), compared to their unlabeled counterparts. **(c)** The functionality of the Lefty construct was assessed by its ability to cause Nodal loss-of-function phenotypes. **(d**,**e)** Schematic of the CRISPR strategy^22^ at the *squint* locus. A target site (cyan) close to the desired insertion site (yellow) is targeted by the Cas9 endonuclease (scissor symbols). (e) After cutting, homology-directed repair (HDR) inserts the donor construct by recognizing its homology flanks (HF). The target site in the donor construct (gray) has been mutated such that it is not targeted itself, but is translated to the same amino acids. **(f)** Sequence confirmation of the *squint-sfGFP* locus. The *squint* locus was PCR-amplified from genomic DNA of adult F2 *squint-sfGFP* fish using the primers GCTGATTCTGTGACCAGCCT + TGCCGCTCTACATGATGCAG and validated by Sanger sequencing. Alignments of sequencing results (red arrows) confirmed the precise integration of sfGFP and the integrity of the *squint* locus, including the homology flank boundary regions (insets). The scale bars are 100 μm in (b) and 300 μm in (c).

**Supplementary Figure 3.**
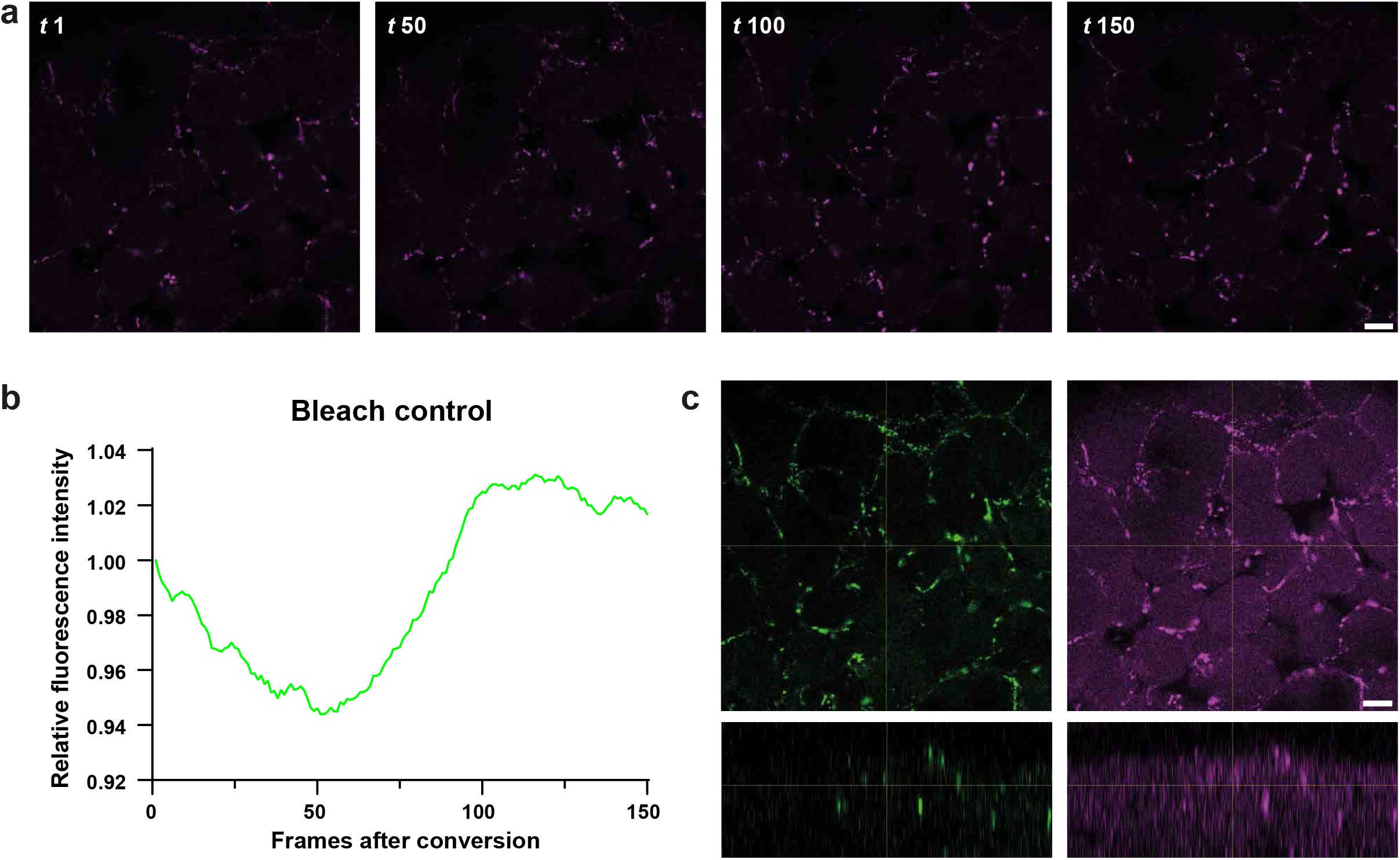
FDAP controls. **(a)** To rule out that bleaching of the fluorescence signal contributes to the apparent protein decay, we imaged Squint-Dendra2 directly after photoconversion for 150 frames at 10-second intervals. The signal maintains the same intensity from frame 1 to frame 150. **(b)** Measurements of fluorescence intensity normalized to timepoint 1. Visible fluctuations (maximum 6%) are due to cell movements. **(c)** z-stack of a representative Squint-Dendra2 embryo showing homogeneous signal conversion along the z axis. Left panels show the unconverted Dendra2 and right panels the converted one. The upper row shows the xy plane, the lower one the orthogonal view in xz. Scale bar: 20 µm, experimental temperature: 28°C.

**Supplementary Figure 4.**
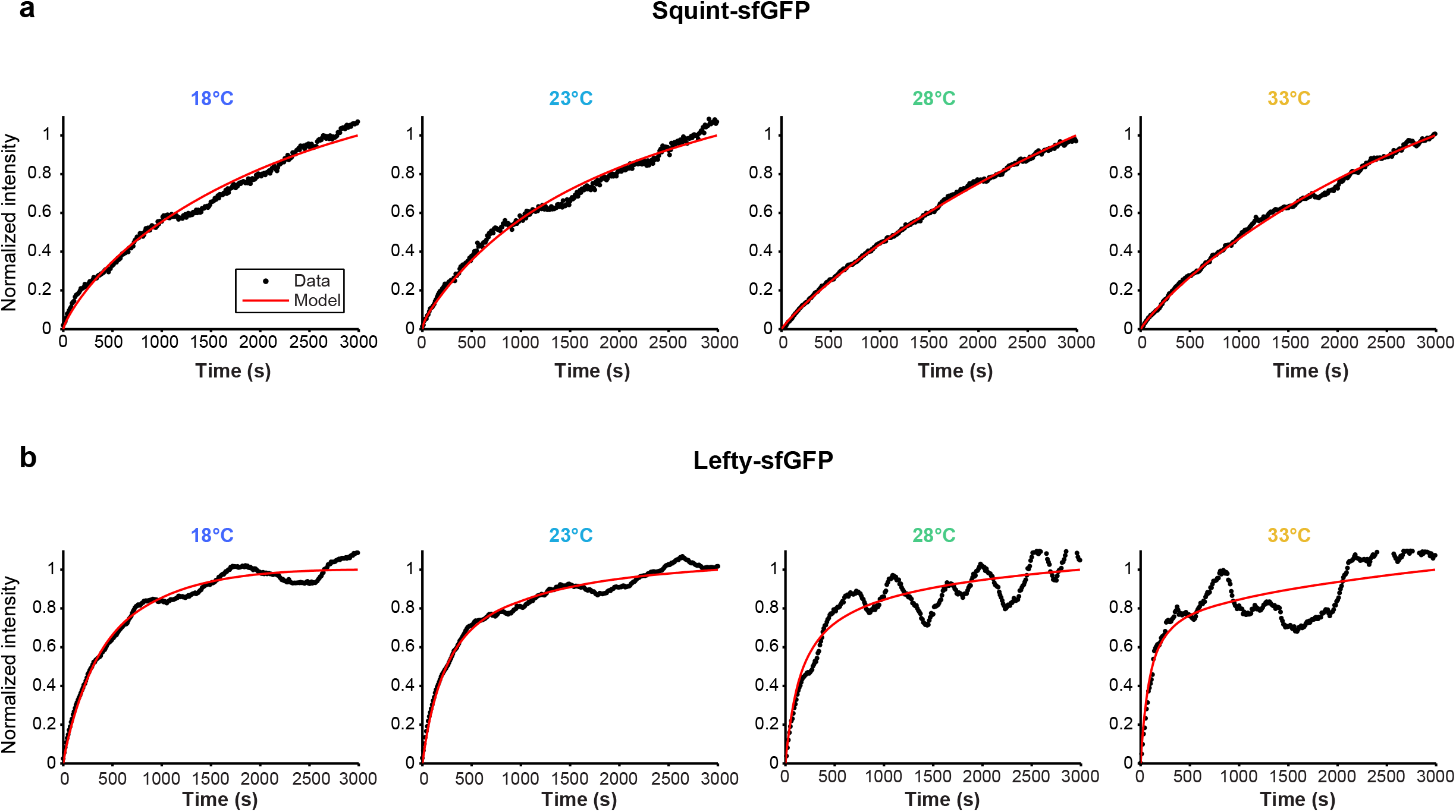
FRAP curves and model fits at different temperatures for Squint and Lefty. Representative normalized fluorescence recovery curves are shown for Squint-sfGFP **(a)** and Lefty-sfGFP **(b)** in medaka embryos at 18°C, 23°C, 28°C, and 33°C (left to right). Black points show the measured recovery and red lines show the fitted model over the displayed 3,000 s interval. Recovery curves were used to determine the diffusion coefficients summarized in Fig. 3c,d.

**Supplementary Figure 5.**
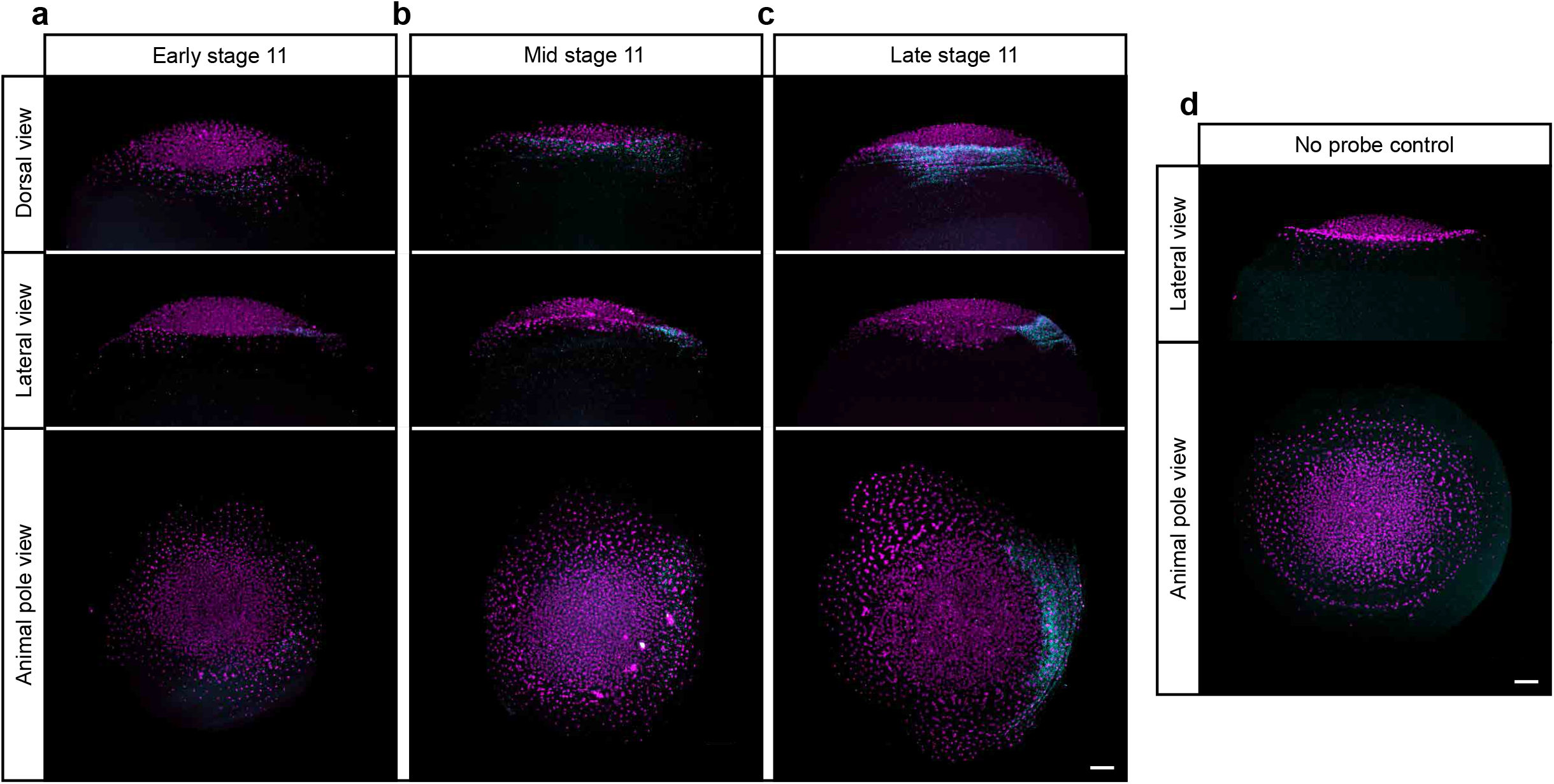
*squint* expression and *in situ* hybridization controls. **(a-c)** Fluorescence *in situ* hybridization shows *squint* expression (cyan) in three representative medaka embryos at progressively later phases of stage 11. The top row shows dorsal views, the middle row lateral views with dorsal to the right, and the bottom row animal-pole views with dorsal to the right. All nuclei were stained with DRAQ7 (magenta). **(d)** *In situ* hybridization control without the *squint*-DIG probe showing the background signal in a stage 11 medaka embryo caused by the anti-DIG-POD antibody in combination with the tyramide signal amplification. Scale bar: 100 µm.

**Supplementary Figure 6.**
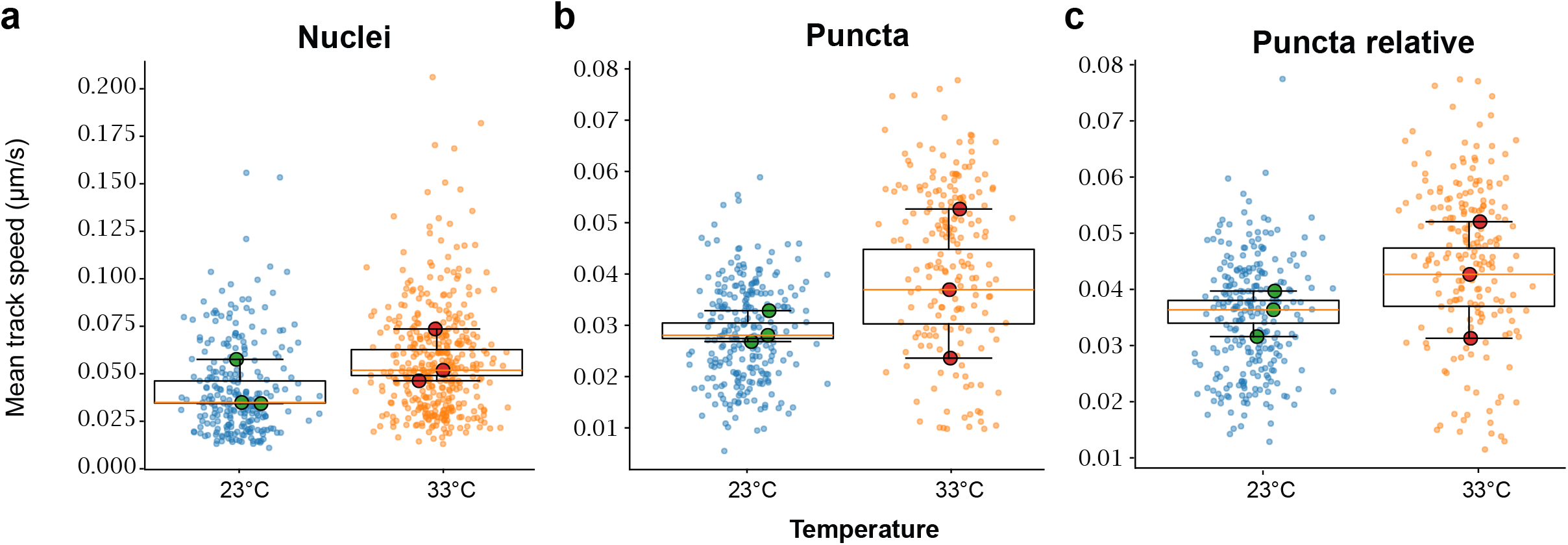
Puncta tracking. (a-c) Mean track speeds of nuclei (a), Squint-sfGFP puncta (b), and Squint-sfGFP puncta after subtraction of the local nuclear velocity field (c). Velocities were calculated from frame-to-frame displacements in three-dimensional light-sheet recordings. Blue and orange points show individual track-speed measurements at 23°C and 33°C, respectively, while large symbols show the mean track speed of single embryos (biological replicates). Panel (c) quantifies residual punctum motion relative to surrounding tissue.

## Supplementary Movie Legends

**Supplementary Movie 1** | **Medaka development at different temperatures**. Brightfield time-lapse recordings of medaka embryos maintained at the indicated temperatures, starting at the one-cell stage (Iwamatsu stage 1). Images were acquired every 2 minutes over 24 hours and are displayed at 20 frames per second. Time indicates elapsed time since the start of imaging. Scale bar: 500 µm.

**Supplementary Movie 2** | **Zebrafish and medaka development over 24 h**. Brightfield time-lapse recordings of zebrafish (left column) and medaka (right column), with two embryos shown for each species. The recordings illustrate the faster progression of zebrafish through gastrulation and embryonic axis elongation compared to medaka. Timestamps indicate elapsed recording time in hours:minutes. Scale bar: 500 μm.

**Supplementary Movie 3** | **Zebrafish and medaka development over 72 h**. Brightfield time-lapse recordings of a zebrafish embryo (left) and a medaka embryo (right), illustrating their different developmental tempi over three days. The sequence follows early embryogenesis, body-axis formation, and subsequent morphological development. Timestamps indicate elapsed recording time in hours:minutes. Scale bar: 500 μm.

**Supplementary Movie 4** | **Nodal inhibition reduces *tbxta:Venus* (*bra:Venus*) reporter expression**. Time-lapse imaging of *tbxta:Venus* medaka embryos^46^ under control conditions or after treatment with 2 or 3 μM Nodal inhibitor starting at Iwamatsu stage 8. Rows show brightfield images (top), Venus fluorescence in grayscale (middle), and merged images with fluorescence in green (bottom). The inhibitor-treated groups show reduced reporter fluorescence during development. Images were acquired every 6 minutes over 24 hours. Scale bar: 500 µm. Playback: 10 frames/s.

**Supplementary Movie 5** | **FDAP bleaching control**. A medaka embryo injected with Squint– Dendra2-encoding mRNA (50 ng/µl) was imaged at 28°C immediately after photoconversion. Images were acquired every 10 seconds for 150 frames using the same acquisition settings as in the standard FDAP experiments, except for the frame interval, and are displayed at 10 frames per second. No appreciable loss of fluorescence was observed despite the increased number of acquisitions compared with the 30 frames used in standard FDAP measurements, indicating that acquisition-induced photobleaching contributes minimally to the measured fluorescence decay. Quantification is shown in Supplementary Fig. 3. Scale bar: 20 µm.

**Supplementary Movie 6** | **FRAP bleaching control**. Uniform bleaching across the imaging depth during FRAP. Maximum-intensity projection of a z-stack of a medaka embryo injected with mRNA encoding cytosolic GFP (50 ng/µl), showing uniform bleaching across the imaged z-range. Scale bar: 40 µm.

**Supplementary Movie 7** | **Squint-sfGFP knock-in**. Time-lapse light-sheet movie recorded every 2 minutes for 12 hours at 28°C. Magenta signal shows H2A-mCherry (50 ng/µl mRNA injected at the one-cell stage) and green the endogenous Squint signal. Squint is first visible in puncta at the blastoderm margin followed by diffuse signal in the periderm. Dorsal view. Scale bar: 50 µm. Playback: 25 frames/s.

**Supplementary Movie 8** | **Epiboly at different temperatures**. Light-sheet time-lapse imaging of Squint-sfGFP knock-in embryos at 18°C and 28°C. Images were recorded every 2 minutes for 2 hours. Magenta signal shows H2A-mCherry (50 ng/µl mRNA injection at 1-cell stage). Nuclei at the margin have been tracked to estimate epiboly speed. Scale bar: 200 µm. Playback: 30 frames/s.

**Supplementary Movie 9** | **Squint-sfGFP at different temperatures**. Light-sheet time-lapse imaging of Squint-sfGFP knock-in embryos at 18°C and 28°C. Images were recorded every 2 minutes. Magenta signal shows H2A-mCherry and green the endogenous Squint signal. Scale bar: 50 µm. Playback: 25 frames/s.

**Supplementary Movie 10** | **Moving-boundary reaction-dominated model of Nodal transport on a growing epiblast domain across developmental temperatures**. Simulated concentration profiles at 18°C (blue), 23°C (cyan), 28°C (green), and 33°C (yellow) are shown as functions of distance from the margin. The model includes production at the margin, diffusion, first-order clearance, and dilution by tissue growth. Production, clearance, and growth scale with a common Arrhenius factor (*E*_*a*_ = 77 kJ/mol), whereas diffusion follows the Einstein-Stokes relation with water-viscosity correction. The displayed time is expressed as a fraction of the temperature-specific simulation duration, allowing comparison at equivalent developmental intervals. The black dashed line indicates the opposite tissue boundary. Concentration is initially zero, and the domain expands from 300 μm as the gradients form.

**Supplementary Movie 11** | **Mathematical model – controls at 18°C**. From left to right, the panels show pure stretching of an initially uniform concentration without production or clearance (*D* = 10^−6^ μm^2^/s, *k* = 0, *j* = 0, *G* = 0.01803 μm/s), production without diffusion or clearance (*D* = 10^−6^ μm^2^/s, *k* = 0, *j* = 3.82 × 10^−5^ mol/(μm^2^·s), *G* = 0.01803 μm/s), production with high diffusion but no clearance (*D* = 10^6^ μm^2^/s, *k* = 0, *j* = 3.82 × 10^−5^ mol/(μm^2^·s), *G* = 0.01803 μm/s), and production with high diffusion and clearance (*D* = 10^6^ μm^2^/s, *k* = 3.82 × 10^−5^/s, *j* = 3.82 × 10^−5^ mol/(μm^2^·s), *G* = 0.01803 μm/s). Concentration *c* is plotted against distance from the margin as the domain expands from an initial length of 300 μm. The pure-stretching control starts at a uniform concentration of 1, and the production controls start at zero. The timestamp indicates simulated time in minutes, covering 0 to 518.3 min.

**Supplementary Movie 12** | **Mathematical model – controls at 23°C**. From left to right, the panels show pure stretching of an initially uniform concentration without production or clearance (*D* = 10^−6^ μm^2^/s, *k* = 0, *j* = 0, *G* = 0.03087 μm/s), production without diffusion or clearance (*D* = 10^−6^ μm^2^/s, *k* = 0, *j* = 6.54 × 10^−5^ mol/(μm^2^·s), *G* = 0.03087 μm/s), production with high diffusion but no clearance (*D* = 10^6^ μm^2^/s, *k* = 0, *j* = 6.54 × 10^−5^ mol/(μm^2^·s), *G* = 0.03087 μm/s), and production with high diffusion and clearance (*D* = 10^6^ μm^2^/s, *k* = 6.54 × 10^−5^/s, *j* = 6.54 × 10^−5^ mol/(μm^2^·s), *G* = 0.03087 μm/s). Concentration *c* is plotted against distance from the margin as the domain expands from an initial length of 300 μm. The pure-stretching control starts at a uniform concentration of 1, and the production controls start at zero. The timestamp indicates simulated time in minutes, covering 0 to 302.7 min.

**Supplementary Movie 13** | **Mathematical model – controls at 28°C**. From left to right, the panels show pure stretching of an initially uniform concentration without production or clearance (*D* = 10^−6^ μm^2^/s, *k* = 0, *j* = 0, *G* = 0.05192 μm/s), production without diffusion or clearance (*D* = 10^−6^ μm^2^/s, *k* = 0, *j* = 1.1 × 10^−4^ mol/(μm^2^·s), *G* = 0.05192 μm/s), production with high diffusion but no clearance (*D* = 10^6^ μm^2^/s, *k* = 0, *j* = 1.1 × 10^−4^ mol/(μm^2^·s), *G* = 0.05192 μm/s), and production with high diffusion and clearance (*D* = 10^6^ μm^2^/s, *k* = 1.1 × 10^−4^/s, *j* = 1.1 × 10^−4^ mol/(μm^2^·s), *G* = 0.05192 μm/s). Concentration *c* is plotted against distance from the margin as the domain expands from an initial length of 300 μm. The pure-stretching control starts at a uniform concentration of 1, and the production controls start at zero. The timestamp indicates simulated time in minutes, covering 0 to 180.0 min.

**Supplementary Movie 14** | **Mathematical model – controls at 33°C**. From left to right, the panels show pure stretching of an initially uniform concentration without production or clearance (*D* = 10^−6^ μm^2^/s, *k* = 0, *j* = 0, *G* = 0.08584 μm/s), production without diffusion or clearance (*D* = 10^−6^ μm^2^/s, *k* = 0, *j* = 1.82 × 10^−4^ mol/(μm^2^·s), *G* = 0.08584 μm/s), production with high diffusion but no clearance (*D* = 10^6^ μm^2^/s, *k* = 0, *j* = 1.82 × 10^−4^ mol/(μm^2^·s), *G* = 0.08584 μm/s), and production with high diffusion and clearance (*D* = 10^6^ μm^2^/s, *k* = 1.82 × 10^−4^/s, *j* = 1.82 × 10^−4^ mol/(μm^2^·s), *G* = 0.08584 μm/s). Concentration *c* is plotted against distance from the margin as the domain expands from an initial length of 300 μm. The pure-stretching control starts at a uniform concentration of 1, and the production controls start at zero. The timestamp indicates simulated time in minutes, covering 0 to 108.9 min.

